# FFCM-MRF: An accurate and generalizable cerebrovascular segmentation pipeline for humans and rhesus monkeys based on TOF-MRA

**DOI:** 10.1101/2023.11.07.566142

**Authors:** Yue Cui, Haibin Huang, Jialu Liu, Mingyang Zhao, Chengyi Li, Xinyong Han, Na Luo, Jinquan Gao, Dongming Yan, Chen Zhang, Tianzi Jiang, Shan Yu

## Abstract

**Purpose:** Cerebrovascular segmentation and quantification of vascular morphological features on humans and rhesus monkeys are essential for prevention, diagnosis, and treatment of brain diseases. However, current automated whole-brain vessel segmentation methods are often not generalizable to independent datasets, limiting their usefulness in real-world environments with their heterogeneity in participants, scanners, and species.

**Materials and Methods:** In this study, we proposed an automated, accurate and generalizable segmentation method for magnetic resonance angiography images called FFCM-MRF. This method integrates fast fuzzy c-means clustering and Markov random field optimization using blood vessel shape priors and spatial constraints. We used a total of 123 human and 44 macaque MRA images scanned at 1.5 T, 3 T, and 7 T MRI from 9 datasets to develop and validate the method.

**Results:** The average Dice score coefficients for multiple independent datasets were 69.16-89.63%, with the improvements in FFCM-MRF ranged from 0.16-16.14% compared with state-of-the-art machine learning methods. Quantitative analysis showed that FFCM-MRF can accurately segment major arteries in the Circle of Willis at the base of the brain and smaller distal pial arteries while effectively suppressing noise. Test-retest analysis showed that the model yielded high vascular volume and diameter reliability.

**Conclusions:** Our results demonstrate that the proposed method is highly accurate and reliable and largely independent of variations in field strength, scanner platforms, acquisition parameters, and species. The macaque MRA data and user-friendly open-source toolbox are freely available at OpenNeuro and GitHub to facilitate studies of imaging biomarkers for cerebrovascular and neurodegenerative diseases.

## 1. Introduction

The structural and functional integrity of blood vessels is essential to normal functioning of the brain. The brain relies highly on a complex and well-regulated vascular network to ensure an adequate supply of blood and meet its high demand for neural activity and metabolism. As a result, blood flow deficiencies and pathological changes in cerebral vessels can affect brain function and contribute to cerebrovascular disease, such as stroke, aneurysms, and arteriovenous abnormalities, as well as neurodegenerative disorders, such as Alzheimer’s disease (1). The rhesus monkey (*Macaca mulatta*) is a commonly used animal model due to its genetic, brain structural, and behavioral similarities to humans (2). A comprehensive delineation and reconstruction of the three-dimensional intracranial vessels for humans and nonhuman primates, along with a quantification of vascular morphological features, could provide insights into the impact of vascular changes on brain health. These may be beneficial for the prevention, diagnosis, and treatment of diseases associated with vascular dysfunction and cognitive decline.

Three-dimensional time-of-flight magnetic resonance angiography (TOF-MRA) is a noninvasive imaging technique for intracranial arterial (and some venous) vessels that is widely used in cerebrovascular clinical and scientific research. However, the task of automated segmentation of cerebral vessels is particularly challenging due to the brain’s complex vasculature and spatially sparse, slender, and tubular structures. Deep learning techniques, such as U-Net and Transformer networks, have recently attracted interest in isolating vessels from brain tissues (3–6). For instance, Livne and colleagues used 2D U-Net for the first time to segment brain vessels and outperformed the traditional graph-cuts method (3). The nnU-Net based on 3D U-Net framework proposed by Isensee et al. (4) is currently state-of-the-art for training biomedical image networks. Chen et al. (6) proposed a generative consistency-based semi-supervised (GCS) model based on Transformer structure in neural networks (7) with semantic fusion for cerebrovascular segmentation. However, deep learning requires high-quality ground truth labels obtained via meticulous, voxel-based manual delineation. This procedure is expensive in terms of time and manpower, dependent on highly trained and experienced experts, and prone to inter-expert variability (8). In addition, deep learning models usually rely on large and heterogeneous training sets with high-quality annotations to prevent overfitting and ensure good performance on independent datasets. However, manual labeling of the whole brain’s sophisticated and complex vasculature network is costly and prone to inter-annotator variability. As a result, publicly available high-quality annotated angiography datasets are limited for human. Furthermore, to the best of our knowledge, there are no open access TOF-MRA data and annotations for nonhuman primates. The lack of enough high-quality training datasets limits the applicability of deep models for cerebrovascular segmentation in humans and macaques. In contrast, unsupervised learning methods, such as clustering and finite mixture model-based methods, capture intrinsic patterns of vessels based on the global properties of the intensity distributions in TOF-MRA (9, 10). Such methods are independent of ground truth labels and adapted to relatively small datasets. However, finite mixture models estimate parameters using an expectation maximization algorithm and are prone to parameter drifting due to unbalanced vascular (∼5%) and nonvascular (∼95%) voxels (11). Fuzzy c-means clustering (FCM) is one of the most widely used techniques in the field of medical image segmentation and can provide better results than expectation maximization (12). Fuzzy logic is a multi-valued logic derived from fuzzy set theory that assigns each voxel a membership in each cluster to a certain degree. FCM clustering method was used in previous studies. For example, in the work of Sakellarios et al., active contour model was used for vascular segmentation and FCM was used for categorization of the type of the plaque (13). Other studies have investigated FCM clustering for brain vessel segmentation on TOF-MRA. However, using FCM clustering alone for TOF-MRA vessel segmentation may result in vascular fragments and non-vessel noises because it does not involve spatial continuity and morphology of vessels (14). To address these limitations, researchers have incorporated vessel enhancement techniques, such as Frangi’s filter (14) and Hessian matrix eigenvalues (15) into the objective function of FCM. However, vessel-enhanced images introduce excessive pseudo vessels, and FCM cannot model spatial neighborhood relationships for continuity, thus these methods generally result in inaccurate vessel segmentation. In contrast, the integration of Markov random field (MRF) with the statistical model allows for spatial similarity constraints among neighboring voxels and tubular structure priors through improved vessel enhancement (16), thereby suppressing non-vessel noises while enhancing the continuity of the segmented vessels. Therefore, we creatively designed a MRF energy function after the FCM segmentation stage with vessel shape and local characteristics constraints in a three-dimensional neighborhood system.

Vessels are spatially heterogeneously distributed across the brain to meet distinct energy and neural interaction requirements (17, 18). Various vascular features including volume, length, and diameter based on a three-dimensional segmentation of vessels provide qualitative mapping and analyses of morphological characteristics, thereby enabling a better understanding of brain structural and functional organization. Most studies have focused on the location and morphology of the Circle of Willis (19, 20) and more distal arteries (17) in humans. Although vascular patterns have become available through human studies, the information about the segmentation, extraction, and quantification of cerebral vessels in rhesus monkeys is still quite limited. There is also a lack of rhesus monkey angiography datasets to promote progress in comparative neuroscience and translational medicine.

In the present study, we developed a novel fast fuzzy c-means clustering with MRF framework (FFCM-MRF) for cerebrovascular segmentation on TOF-MRA. Tubular-shaped priors of blood vessels and spatial neighborhood information were optimized by MRF to remove noise and improve segmentation accuracy and spatial continuity. The proposed method was validated on multiple independent human and macaque datasets and compared with state-of-the-art methods. Vascular volume density and diameter features were then extracted and analyzed for humans and macaques. We also developed a user-friendly MATLAB toolbox for FFCM-MRF vessel segmentation and the analysis of TOF-MRA images.

## 2. Materials and methods

### 2.1 MRA datasets

A total of 123 human and 44 macaque MRA images scanned at 3 T and 7 T MRI from 9 datasets was used to develop and validate FFCM-MRF. Human MRA data were taken from the publicly available MIDAS, BraVa, CHUV, ADAM, Forrest, and Midnight Scan Club (MSC) datasets. 36 Macaque MRA images were acquired from a 3 T Siemens Prisma scanner in Wuhan (Maca-WH), 4 were acquired from a 3 T Siemens Prisma scanner from the Institute of Biophysics, Chinese Academy of Sciences in Beijing (Maca-BJ), and 4 were obtained from a 7 T MAGNETOM Terra scanner (Maca-7T) at the same institute. See Tables 1 and 2 for subject details and references for acquisition parameters. Details of these datasets and ground truth labels refer to Supplementary Methods. This study was approved by the Institutional Review Board/Ethics Committee of Chinese Academy of Sciences Institute of Automation.

**Table 1.** Subject demographics ^a^.

|  | Dataset | Subject number | Age (years) | Sex (F/M) |
| --- | --- | --- | --- | --- |
| Humans | MIDAS | 54 | 38.1 (12.3) | 23/31 |
|  | BraVa | 5 | 28.8 (8.9) | 1/4 |
|  | CHUV | 5 | - | - |
|  | ADAM | 5 | - | - |
|  | Forrest | 14 | 20 ~ 35 | 4/9 |
|  | MSC | 10 (40 scans) | 29.1 (3.2) | 5/5 |
| Macaques | Maca-WH | 36 | 9.4 (4.0) | 36/0 |
|  | Maca-BJ | 4 | 4.0 (0.6) | 4/0 |
|  | Maca-7T | 4 | 4.5 (1.0) | 4/0 |

<sup>a</sup> Age and sex information are not available for the ADAM and CHUV datasets. Values are mean (SD) for age except in the Forrest dataset, which uses a range.

**Table 2.**
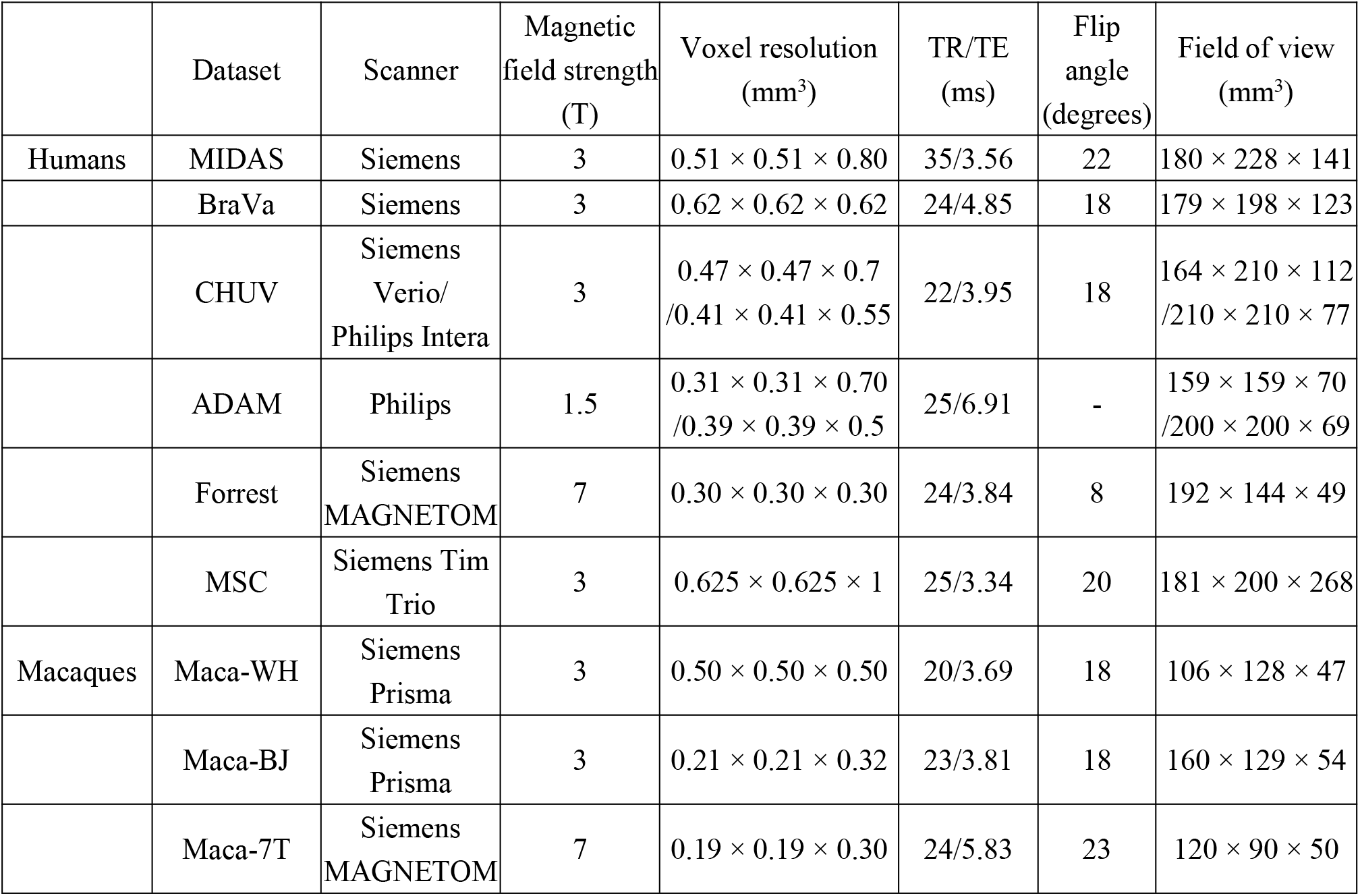
TOF-MRA acquisition parameters.

|  | Dataset | Scanner | Magnetic field strength (T) | Voxel resolution (mm<sup>3</sup>) | TR/TE (ms) | Flip angle (degrees) | Field of view (mm<sup>3</sup>) |
| --- | --- | --- | --- | --- | --- | --- | --- |
| Humans | MIDAS | Siemens | 3 | $0.51 \times 0.51 \times 0.80$ | 35/3.56 | 22 | $180 \times 228 \times 141$ |
| | BraVa | Siemens | 3 | $0.62 \times 0.62 \times 0.62$ | 24/4.85 | 18 | $179 \times 198 \times 123$ |
| | CHUV | Siemens Verio/<br>Philips Intera | 3 | $0.47 \times 0.47 \times 0.7$<br>/$0.41 \times 0.41 \times 0.55$ | 22/3.95 | 18 | $164 \times 210 \times 112$<br>/$210 \times 210 \times 77$ |
| | ADAM | Philips | 1.5 | $0.31 \times 0.31 \times 0.70$<br>/$0.39 \times 0.39 \times 0.5$ | 25/6.91 | - | $159 \times 159 \times 70$<br>/$200 \times 200 \times 69$ |
| | Forrest | Siemens MAGNETOM | 7 | $0.30 \times 0.30 \times 0.30$ | 24/3.84 | 8 | $192 \times 144 \times 49$ |
| | MSC | Siemens Tim Trio | 3 | $0.625 \times 0.625 \times 1$ | 25/3.34 | 20 | $181 \times 200 \times 268$ |
| Macaques | Maca-WH | Siemens Prisma | 3 | $0.50 \times 0.50 \times 0.50$ | 20/3.69 | 18 | $106 \times 128 \times 47$ |
| | Maca-BJ | Siemens Prisma | 3 | $0.21 \times 0.21 \times 0.32$ | 23/3.81 | 18 | $160 \times 129 \times 54$ |
| | Maca-7T | Siemens MAGNETOM | 7 | $0.19 \times 0.19 \times 0.30$ | 24/5.83 | 23 | $120 \times 90 \times 50$ |

### 2.2 Automated cerebrovascular segmentation procedure

The automated cerebrovascular segmentation procedure consisted of three stages: MRA pre-processing, rudimentary segmentation using fast fuzzy c-means clustering, and a neighborhood constraint energy function for maximum a posteriori information with MRF. These stages are illustrated in Figure 1 and described in detail below.

**Figure 1.**
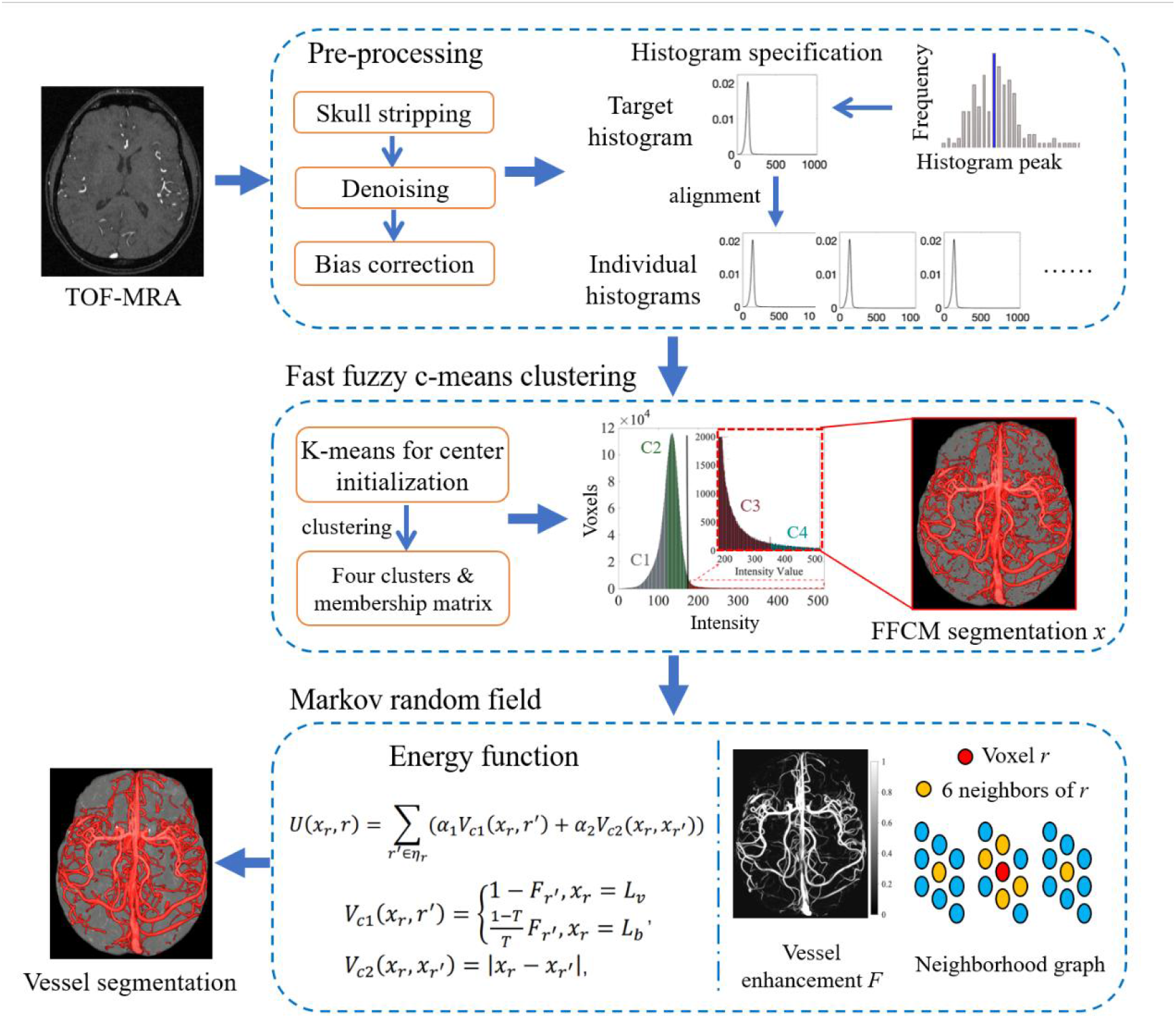
Flowchart of the cerebrovascular segmentation using fast fuzzy c-means clustering combined with Markov random field refinement by spatial and shape constraints.

#### 2.2.1 MRA pre-processing

The MRA images were skull stripped, denoised and bias field corrected. To reduce the intensity variability across subjects, we used histogram specification to improve the robustness of the segmentation algorithm (Supplementary Methods).

#### 2.2.2 Rudimentary segmentation using fast fuzzy c-means clustering

In the present study, fast FCM was used for rudimentary cerebrovascular segmentation. FCM is a data clustering technique in which voxels are grouped into c clusters using fuzzy memberships. The fast FCM calculates the membership matrix and the cluster centers based on the histogram rather than on individual voxels to decrease the execution time of the conventional FCM (21). As a result, the membership calculation was dramatically reduced from an average of more than 6,500k to 0.5 - 1k values. Let *Y = {y_1_,y_2_,*⋯*,y_S_}* denote an MRA image with *S* histogram bins to be partitioned into *C(2*≤ *C* ≤ *S)*clusters, where *y_i_* represents the intensity of voxels in the histogram bin *i* ∈ *1,*…*,S*, and *H_i_* is the frequency of the intensity level at *i*.

First, histogram-based k-means clustering was used to cluster the target histogram, and the centers of clusters *Z_k_*, *k = 1,*⋯*,C,* were used to initialize a fast FCM algorithm for each subject. The membership matrix was then calculated as follows:

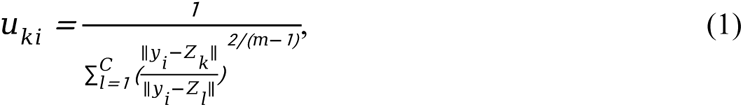

where *m (m >1)*is the fuzzy factor, controlling the fuzziness or probability of the resulting partition. The centers of clusters were calculated by:

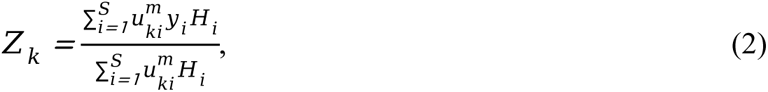

The algorithm iteratively updates the membership values *u_ki_* and cluster centers *Z_k_,(k = 1,2,⋯,C)*according to (1) and (2), until convergence with max 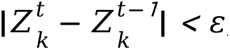, where ε is a user-defined threshold. At the end of the iterations, a membership matrix with a size of *S* × *C* was obtained, which contains the membership probability for voxels at each intensity level belonging to each cluster.

Note that when ||*y_i_* − *Z_k_*||*=0,* there may arise a singularity problem since the membership *u_ki_ =1*, which leads to the failure of the subsequent MRF algorithm. To solve this problem, the membership matrix was clipped within the range of 0 and 1 by:

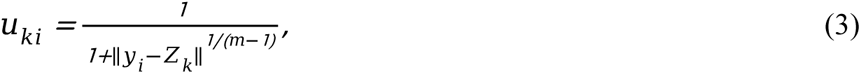

Finally, each voxel r in TOF-MRA can be labeled with 1 (vessel) or 0 (background) as follows:

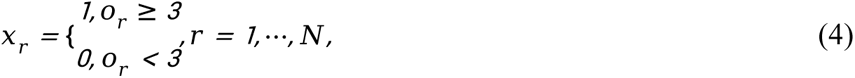

where *x_r_* represents the label of current voxel *r*, *o_r_* is the highest membership value defined by:

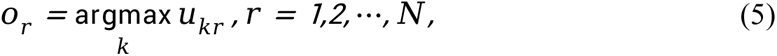

where *u_kr_* denotes the corresponding membership of voxel *r* with cluster *k* (identical with *u_ki_*, i.e., voxel *r* within histogram bin *i*), and *N* is the number of voxels in the intracranial brain tissue.

Experimentally, we compared regions with vessels for three- and four-cluster FCM, and found that when the cluster number *C = 4*, voxels in the first and second clusters correspond to cerebrospinal fluid and brain tissues, respectively, while voxels in the third and fourth clusters corresponded to vessels and were a rudimentary segmentation of the vessels (Supplementary Figure 1).

To further enhance the robustness of the above developed FFCM algorithm for brain vessel segmentation, spatial constraints between neighborhood voxels were modeled by a MRF energy function to reduce noise and increase the integrity of the segmented vessels. The energy function was defined as follows:

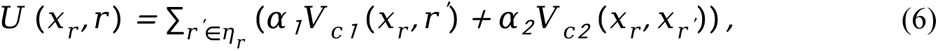

where *x_r_* represents the label of the current voxel *r*, *x_r_’* represents the label of one of the adjacent voxels of *r*, and η*_r_* is the set of six spatial neighbors of *r*. The initial setting of *x* is FCM segmentation (0 or 1), and *x* is then iteratively updated by MRF. Both α*_1_* and α*_2_* were set to 0.5 in the present study (9). *V_c1_* and *V_c2_* are defined as follows:

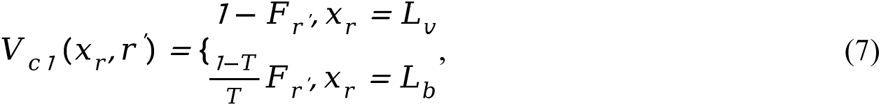

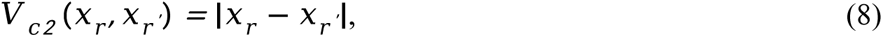

where *L_v_* represents vessel voxels, and *L_b_* represents non-vessel voxels. *F* (*0*≤ *F* ≤ *1* is the Hessian-based multi-scale filtering response for vessel enhancement, which can effectively highlight tubular structure and suppress non-vascular tissues (16). *T* is the threshold that defines the vessel voxels in *F* and can be calculated by

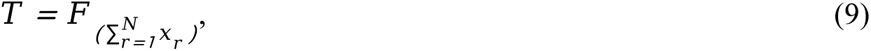

where *F_(n)_* represents the *n*^th^ highest value among all the voxel response values in *F*.

Based on the membership matrix and class prior probability, we adopted the iterated conditional modes to iteratively update the MRF. Based on the equivalence of MRF and Gibbs, class prior probability can be calculated by:

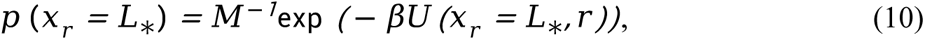

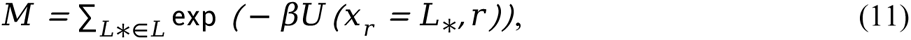

where the parameter β describes the strength of the interaction between pair-wise neighboring voxels. The vascular posterior probability and segmentation criteria can be obtained by:

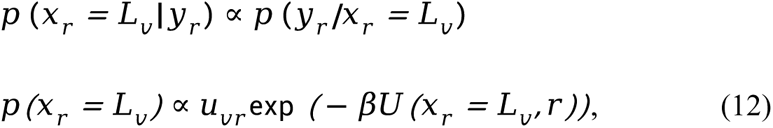

Likewise,

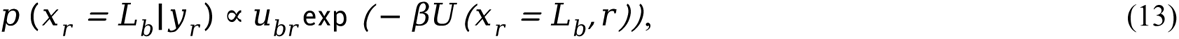

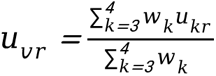 and 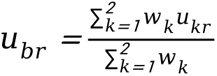, where *w_k_* denotes the proportion of a voxel number in cluster *k* that was determined in the FCM segmentation.

If a voxel meets *p* (*x_r_=L_v_* |*y_r_*) *> p* (*x_r_ = L_b_*|*y_r_*), it is considered a vascular voxel and thus updates the labels *L_v_* and *L_b_*. The MRF optimization is applied iteratively to update the labels until it reaches the maximum iteration (i.e., itermax).

We randomly selected 50% of the participants from MIDAS to tune the parameters *m* and *β* by maximizing the Dice score coefficient (DSC), which evaluates the overlap index between segmentation and ground truth, and is defined as DSC *=2*TP/*(2*TP+FP+FN), where TP is the number of true positive voxels, FP is the number of false positive voxels and FN is the number of false negative voxels. Optimal parameters *m* and *β* were then applied to the whole MIDAS, as well as to the independent human (BraVa, CHUV, ADAM and Forrest) and macaque (Maca-WH, Maca-BJ and Maca-7T) datasets.

### 2.3 FFCM-MRF quantitative evaluation

The complete evaluation and validation of FFCM-MRF were based on the results from qualitative, quantitative, and test-retest reliability assessments. For the qualitative and quantitative analyses, FFCM-MRF was compared with other state-of-the-art methods based on the consistency between the segmentation and the ground truth. The DSC, precision score, recall score, average Hausdorff distance (AHD), 95% Hausdorff distance (95HD), and islands metrics were evaluated on the human and macaque datasets. The precision score measures the proportion of correctly segmented vascular voxels relative to the total number of segmented voxels, i.e., Precision *=*TP/(TP+FP). Recall measures the proportion of correctly segmented vascular voxels relative to the number of voxels in the ground truth, i.e., Recall *=*TP/(TP+FN). AHD and 95HD were used to evaluate the distance between the segmentation and ground truth, see equations in the Supplementary files. In addition, the number of islands was used to indicate the connectivity of the cerebrovascular system. A low number of islands means a more connected vascular structure.

### 2.4 Comparisons with other state-of-the-art models

FFCM-MRF was compared with two state-of-the-art deep neural network models, nnU-Net (4) and GCS (6), as well as with a statistical model, GMM-MRF (9). The TOF-MRA images were pre-processed using skull stripping, denoising, bias field correction, and histogram specification before input to models. Details of state-of-the-art methods are in the Supplementary Methods.

### 2.5 Test-retest reliability

For the reliability assessment, FFCM-MRF was applied to the human MIDNIGHT MSC database and rhesus monkey test-retest data. MSC database contains 10 participants with 4 MRAs for each participant. Rhesus monkey test-retest data consists of macaques M14 and M28. M14 was scanned twice using a 3 T Siemens scanner, and M28 was scanned using 3 T and 7 T Siemens scanners. In each image, the vascular volume density and average diameter of each region of interest (ROI) were computed according to the human (22) and macaque (23) Brainnetome atlases, which included 210 and 248 ROIs for human and macaque cortices, respectively. These ROIs were obtained by anatomical landmarks and structural connectivity profiles by diffusion MRI. The diameter was calculated based on resampling the vascular segmentation at equal intervals for anisotropic images, and the centerline of the vessels was obtained by skeletonizing the vascular segmentation. Resampling the vascular segmentation at equal intervals was performed using the ResampleImageFilter class from SimpleITK library in Python. The diameters were the maximum possible sphere fitting for each vessel skeleton voxel (24).

An intra-class correlation coefficient (ICC) was used to assess the reliability of the vascular volume density and diameter across scans in the MSC dataset. ICC estimates and their 95% confidence intervals were computed based on a single rater/measurement, absolute agreement, two-way mixed effects model (ICC(3,1)). This was conducted with Python 3.7.3 using the Pingouin package (v.0.3.11, https://pingouin-stats.org/build/html/index.html).

### 2.6 FFCM-MRF application

The group-averaged ROI-based vascular volume density and diameters were calculated for the subjects from the MIDAS (*n* = 54) and Maca-WH (*n* = 36) databases based on the human and macaque Brainnetome atlases for cerebrovascular evolutionary comparisons. Specifically, each native TOF-MRA image was first linearly then nonlinearly warped to the structural template in atlas space. The derived transformation parameters were then used to warp vascular density and diameter measurements from native space to atlas space. The registration procedures were performed using Aladin and F3D tools from the NiftyReg package (https://sourceforge.net/projects/niftyreg/). The measurements in each volume-based ROI were averaged across subjects and then *z* scored across cortex. The normalized measurements were assigned to surface-based atlas and visualized using Connectome Workbench (https://www.humanconnectome.org/software/connectome-workbench). In addition, voxelwise feature maps were computed inside the bounding boxes of 15-cube human and 8-cube macaques, with each brain voxel as the center of the bounding box. The features within a bounding box were allocated to the voxel. Volume-based vascular density and diameter measurements were mapped to individual surfaces. Note the individual surfaces were constructed with native T1- and T2-weighted images using HCP pipeline for humans (https://github.com/Washington-University/HCPpipelines) and NHP pipeline for macaques (https://github.com/Washington-University/NHPPipelines) that used a nonlinear volume registration to align with the MNI coordinate system. The surface-based measurements were averaged across subjects and then *z* scored across cortex.

## 3. Results

### 3.1 Qualitative and quantitative assessment

The hyper-parameters tuned on MIDAS for each method are presented in Supplementary Figure 2. For GMM-MRF, the optimal *β* was 0.1. For FFCM-MRF, the optimal *β* was between 0.001 to 0.01, and *m* was between 1.1 and 1.9; so we set *β* to 0.01 and *m* to 1.7 for all datasets. The complete quantitative results for the various methods are shown in Table 3 for multiple datasets. Specifically, on the MIDAS database, nnU-Net outperformed FFCM-MRF, GCS, and GMM-MRF methods in DSC, Precision, AHD and 95HD. The DSC scores for nnU-Net and FFCM-MRF on MIDAS were 90.96% and 85.25%, respectively. While for independent human and macaque datasets, the FFCM-MRF significantly outperformed nnU-Net, GCS, and GMM-MRF in terms of DSC (paired t test between FFCM-MRF and three models with corrected *p* < .05). Compared with nnU-Net, the improvements in FFCM-MRF ranged from 0.16% - 6.74% across independent datasets. The superiority of FFCM-MRF was replicable with different epochs of nnU-Net (Supplementary Figure 3).

**Table 3.** Cerebrovascular segmentation results from four different methods for each dataset in terms of Dice similarity coefficient (DSC), precision score, recall score, average Hausdorff distance (AHD), 95% Hausdorff distance (95HD), and islands metrics ^a^.

| Datasets | Methods | DSC (%) ↑ | Precision(%) ↑ | Recall (%) ↑ | AHD ↓ | 95HD ↓ | Islands ↓ |
| --- | --- | --- | --- | --- | --- | --- | --- |
| MIDAS | nnU-Net<br>(Isensee et al., 2021) | **90.96 (2.17)** | **93.18 (2.43)** | 88.94 (3.63) | **0.25 (0.10)** | **27.75 (4.53)** | 180.48 (21.57) |
|  | GCS<br>(Chen et al., 2023) | 84.33 (3.16) | 82.99 (7.49) | 86.65 (6.08) | 0.46 (0.17) | 35.29 (6.06) | 459.24 (128.60) |
|  | GMM-MRF<br>(Zhang et al., 2019) | 77.14 (2.70) | 72.56 (4.81) | 82.74 (4.14) | 0.71 (0.22) | 36.11 (6.35) | **103.65 (20.95)** |
|  | FFCM-MRF | 85.25 (3.99) | 82.45 (8.46) | **89.05 (3.74)** | 0.69 (0.33) | 35.99 (6.39) | 133.30 (38.72) |
| BraVa | nnU-Net | 83.06 (2.72) | 88.36 (9.17) | 79.37 (7.10) | 0.56 (0.27) | 38.54 (5.20) | 241.00 (19.67) |
|  | GCS | 83.97 (5.22) | **93.74 (5.09)** | 77.00 (11.02) | **0.39 (0.29)** | 42.42 (10.89) | 210.40 (19.05) |
|  | GMM-MRF | 82.01 (7.34) | 75.01 (13.34) | **92.70 (6.90)** | 1.28 (0.79) | 40.63 (5.01) | **69.60 (26.69)** |
|  | FFCM-MRF | **85.45 (5.64)** | 90.16 (9.14) | 83.16 (13.56) | 0.54 (0.22) | **37.75 (7.60)** | 72.00 (22.90) |
| CHUV | nnU-Net | 86.36 (1.44) | 85.41 (5.94) | 87.89 (5.00) | 0.44 (0.08) | **47.96 (9.57)** | 71.00 (17.68) |
|  | GCS | 82.01 (6.98) | 71.70 (11.91) | **97.19 (1.72)** | 0.95 (0.47) | 62.01 (12.53) | 363.00 (262.63) |
|  | GMM-MRF | 87.04 (4.42) | 87.16 (13.56) | 90.09 (13.47) | 0.44 (0.19) | 57.15 (9.06) | **59.80 (25.19)** |
|  | FFCM-MRF | **89.16 (1.27)** | **92.54 (9.54)** | 87.68 (10.40) | **0.37 (0.17)** | 56.76 (12.36) | 77.60 (49.28) |
| ADAM | nnU-Net | 83.66 (4.57) | 78.88 (10.47) | 90.30 (4.47) | 0.64 (0.10) | 58.01 (7.12) | **73.60 (17.56)** |
|  | GCS | 81.49 (13.83) | 82.37 (24.54) | 87.05 (10.96) | 1.27 (1.79) | 69.29 (14.43) | 259.20 (213.10) |
|  | GMM-MRF | 87.87 (7.82) | 83.15 (16.78) | **95.70 (5.17)** | 0.53 (0.31) | 59.73 (17.56) | 87.60 (11.33) |
|  | FFCM-MRF | **89.63 (5.85)** | **92.11 (10.56)** | 89.04 (12.01) | **0.34 (0.20)** | **52.44 (13.54)** | 76.60 (13.89) |
| Forrest | nnU-Net | 70.32 (7.08) | 87.73 (4.23) | 59.60 (10.26) | 1.91 (0.70) | 71.31 (6.46) | 567.64 (47.29) |
|  | GCS | 70.82 (8.30) | **93.46 (3.08)** | 57.89 (10.67) | 1.14 (0.40) | 56.47 (6.02) | 429.57 (127.78) |
|  | GMM-MRF | 76.44 (6.57) | 81.66 (7.62) | **73.70 (12.44)** | **0.81 (0.30)** | **54.19 (9.04)** | 368.57 (107.45) |
|  | FFCM-MRF | **77.06 (6.81)** | 84.16 (6.58) | 72.73 (12.45) | 0.94 (0.35) | 57.11 (8.40) | **208.79 (50.25)** |
| Maca-WH | nnU-Net | 64.36 (3.62) | 89.04 (4.67) | 50.75 (5.31) | **0.92 (0.18)** | 25.15 (4.72) | 63.44 (14.34) |
|  | GCS | 53.02 (4.52) | **93.45 (3.04)** | 37.21 (4.56) | 1.82 (0.34) | 28.44 (3.54) | 51.75 (14.71) |
|  | GMM-MRF | 65.84 (4.26) | 65.64 (6.36) | **67.08 (8.26)** | 1.93 (0.85) | 30.89 (15.31) | 71.58 (22.15) |
|  | FFCM-MRF | **69.16 (4.23)** | 82.83 (8.10) | 59.85 (4.97) | 1.19 (0.42) | **23.85 (3.49)** | **22.69 (8.34)** |
| Maca-BJ | nnU-Net | 72.81 (3.79) | 65.49 (5.96) | **82.34 (3.05)** | 3.32 (0.73) | 83.31 (5.60) | 201.75 (71.55) |
|  | GCS | 71.33 (2.56) | **84.95 (5.44)** | 61.92 (6.09) | 2.23 (0.51) | 77.37 (6.71) | 176.25 (50.55) |
|  | GMM-MRF | 65.39 (4.28) | 66.20 (6.39) | 65.37 (6.97) | **1.67 (0.46)** | **25.33 (2.88)** | **65.40 (13.40)** |
|  | FFCM-MRF | **72.97 (5.02)** | 66.96 (8.80) | 80.89 (3.19) | 3.07 (0.97) | 80.37 (6.36) | 101.75 (43.34) |
| Maca-7T | nnU-Net | 79.07 (7.88) | 73.12 (14.03) | **87.94 (5.53)** | **0.65 (0.37)** | **50.31 (16.75)** | 211.50 (38.16) |
|  | GCS | 69.87 (12.74) | **88.61 (13.31)** | 62.29 (22.64) | 1.16 (0.45) | 64.43 (27.49) | 260.25 (190.57) |
|  | GMM-MRF | 76.40 (7.47) | 70.54 (14.93) | 86.16 (6.34) | 0.92 (0.62) | 54.02 (18.31) | **124.75 (126.94)** |
|  | FFCM-MRF | **80.24 (8.52)** | 78.71 (16.61) | 83.98 (7.18) | 1.01 (1.13) | 55.50 (13.17) | 128.50 (50.43) |

<sup>a</sup> Values are mean (SD) unless otherwise indicated. Bold denotes the best results from the four different methods.

The detailed qualitative analysis of the segmentation results using state-of-the-art methods on MIDAS are shown in Figure 2. Note that the vasculature annotations were often ignored for distal small arteries, while the proposed FFCM-MRF was superior in detecting vessels that were missed as the ground truth labels (see also Supplementary Figure 4). Qualitative analysis of the segmentation results on new images from the independent datasets showed that the proposed FFCM-MRF is capable of segmenting large blood vessels, including the internal carotid arteries (ICA, Figure 3B) and smaller distal arteries (Figure 3). In addition, our method is more robust to noise than GMM-MRF perhaps because it benefits from the effective Markov optimization procedure.

**Figure 2.**
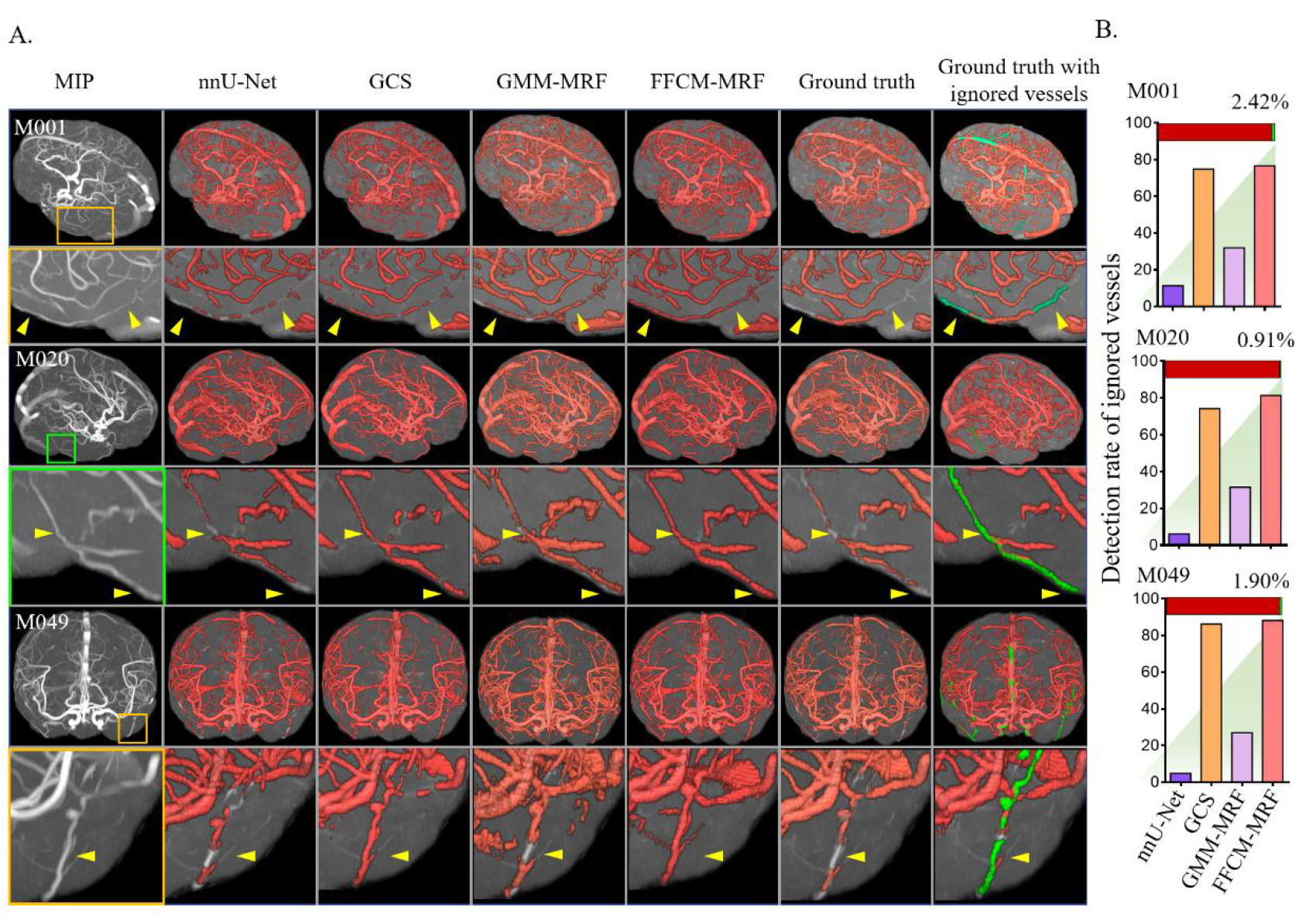
Comparisons of the cerebrovascular segmentation results from the MIDAS dataset using state-of-the-art methods show that FFCM-MRF was superior in detecting ignored vessels. A, Left to right columns correspond to the maximum intensity projection (MIP) views for the TOF-MRA image, the results from nnU-Net, GCS, GMM-MRF, and FFCM-MRF, ground truth, and the ground truth with ignored vessels shown in green. The yellow arrows indicate that FFCM-MRF can detect vessels that were not annotated in the ground truth. Three subjects were meticulously delineated by a trained annotator with the ignored vessels in green, as shown in the rightmost column in A. B, Detection rate of ignored vessels with state-of-the-art methods. Ignored vessels account for a small portion of the whole brain vessels on TOF-MRA, indicated by the green (ignored) and red (ground truth) colors of the horizontal bars in B.

**Figure 3.**
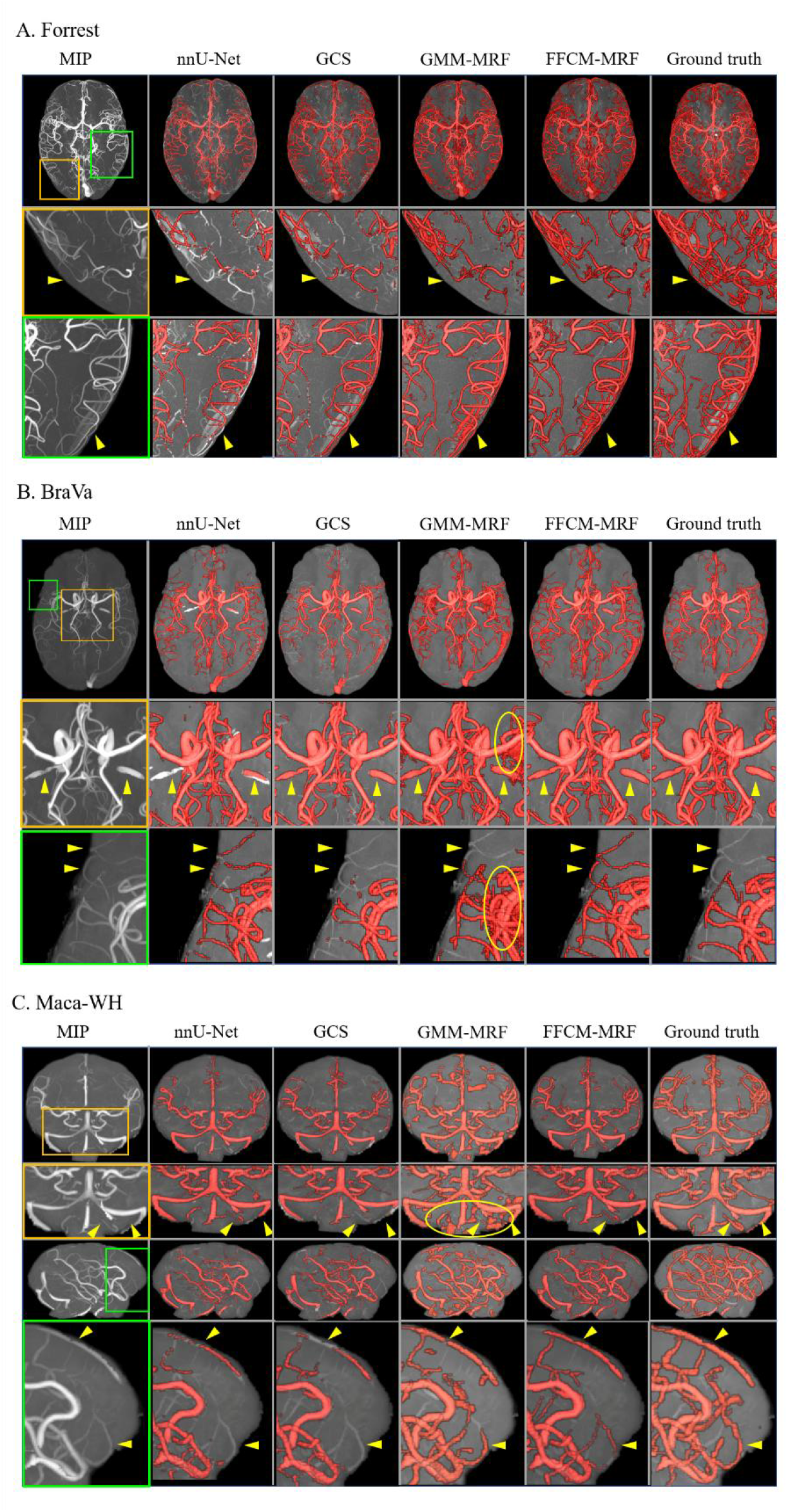
Qualitative comparisons of the cerebrovascular segmentation results from independent datasets using state-of-the-art methods. Left to right columns correspond to a maximum intensity projection (MIP) view for the TOF-MRA image. The results are for nnU-Net, GCS, GMM-MRF, and FFCM-MRF as well as for ground truth. Yellow arrows indicate that deep learning models derived from the MIDAS dataset performed poorly on the Forrest (A), BraVa (B) and Maca-WH (C) datasets for large arteries (B) and distal smaller vessels (A, B, and C). Yellow ellipses show that FFCM-MRF was more robust to noise than GMM-MRF.

Detailed comparisons between nnU-Net and FFCM-MRF showed that nnU-Net often had poorer detection of vessels near the edge of the brain. These included the Circle of Willis (CW) arteries located at the base of the brain and smaller distal arteries that cover the cerebral cortical surface (Figure 4A). Because the CW plays a vital role in brain health and provides an anastomotic connection between the anterior and posterior circulations and between the left and right hemispheres, we labeled the CW arteries using an overlap of an automated CW segmentation pipeline (20) and employing the ground truths and obtained the detection rate (the proportion of segmentation in eight CW arteries) using nnU-Net and FFCM-MRF (Figure 4 and Supplementary Table 2). We observed that the detection results of the CW arteries using FFCM-MRF outperformed nnU-Net for most of the large arteries on MIDAS and independent datasets (Figure 4B). In addition, FFCM-MRF provided ∼22X faster parameter tuning time using CPU than the nnU-Net model training time using GPU. The GPU inference speed for nnU-Net was faster than FFCM-MRF, which uses CPU, while the CPU inference speed for nnU-Net is much slower than FFCM-MRF (∼2X, Supplementary Table 3).

**Figure 4.**
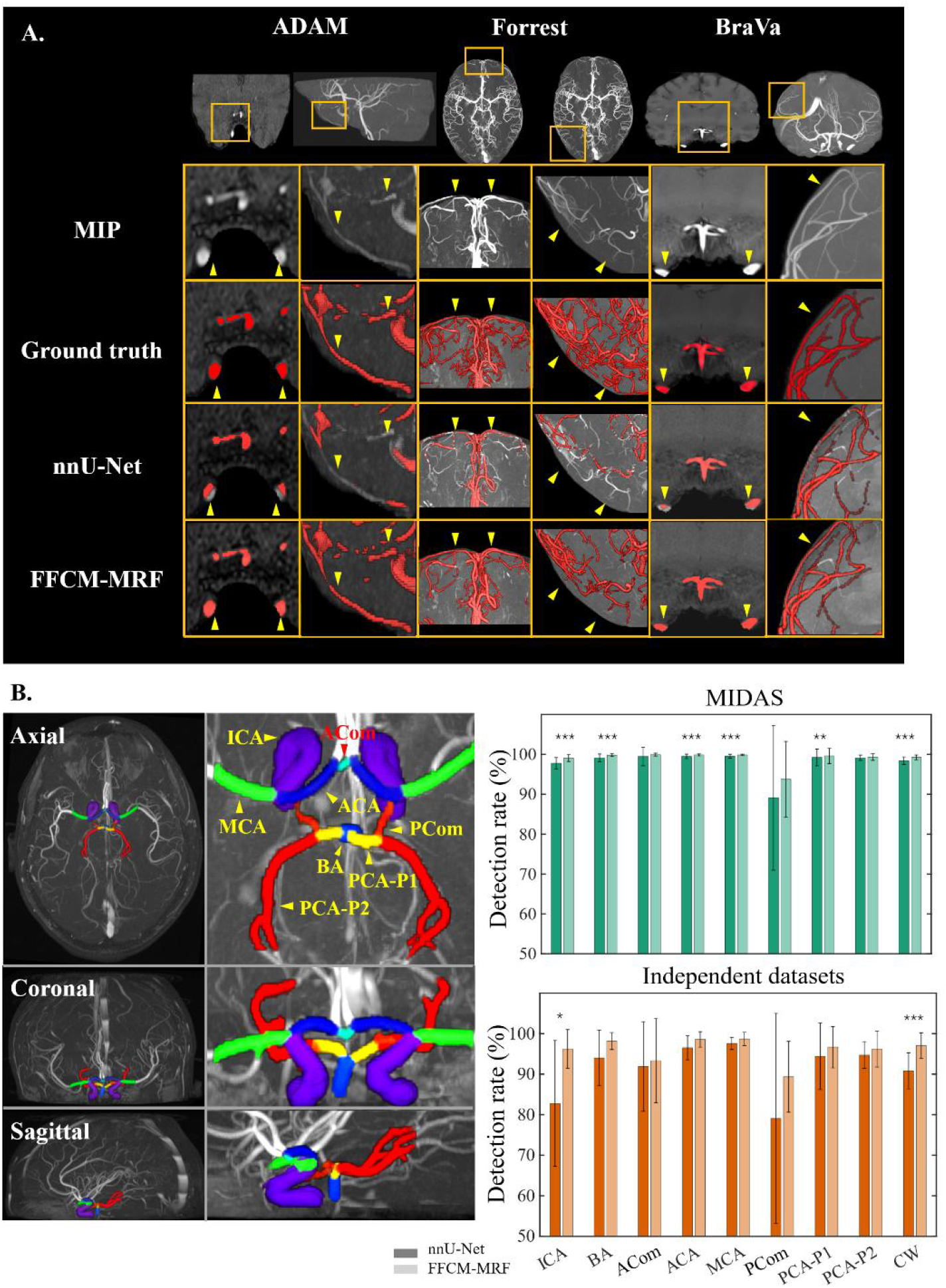
Comparison of segmentation results between nnU-Net and FFCM-MRF. A, Vessels near the edge of the brain/image, including major large vessels at the base of the brain and smaller distal arteries covering the cerebral cortical surface, were not detected by nnU-Net. Note that the ADAM MRA images only partially covered the brain. B, Comparison of nnU-Net and FFCM-MRF in segmenting the circle of Willis (CW). CW arteries were labeled using Dumais et al. (2022). Bar plots showing the proportion of segmented vessels in each CW artery and in the total CW using nnU-Net and FFCM-MRF. The Forrest MRA data only partially covered the CW and were not labeled in this study. ACA, anterior cerebral artery; ACom, anterior communicating artery; BA, basilar artery; ICA, internal carotid artery; MCA, middle cerebral artery; PCA, posterior cerebral artery; PCom, posterior communicating artery. Paired t tests with Bonferroni correction were performed between nnU-Net and FFCM-MRF. ***, corrected *p* < .001; **, corrected *p* < .01; *, corrected *p* < .05.

### 3.2 Test-retest reliability

The reliability of the vascular volume density and average diameter using the human Brainnetome atlas is shown in Figure 5A and Supplementary Table 4. Most regions exhibited good (0.72 - 0.90) to excellent (> 0.90) ICC values across four scans with the exception of the middle temporal gyrus and lateral occipital cortex in volume density. The diameters showed less reliability than the volume densities. Across subjects, the average ICCs were 0.996 and 0.863 for the total brain vascular volume and average within-brain diameter, respectively (Supplementary Figure 5). A comparison of vascular volume density and diameter by the macaque Brainnetome atlas between two 3 T scans from a single subject is shown in Figure 5B. The intra-subject results show that FFCM-MRF provides highly consistent measures of vascular morphology. Figure 5C shows the differences between 3 T and 7 T scans from a macaque. We observed that the vessels were denser from the 7 T MRA than from those detected by the 3 T MRA. The average diameter from 7 T was smaller than those from 3 T, indicating that 7 T MRA can detect more slender vessels.

**Figure 5.**
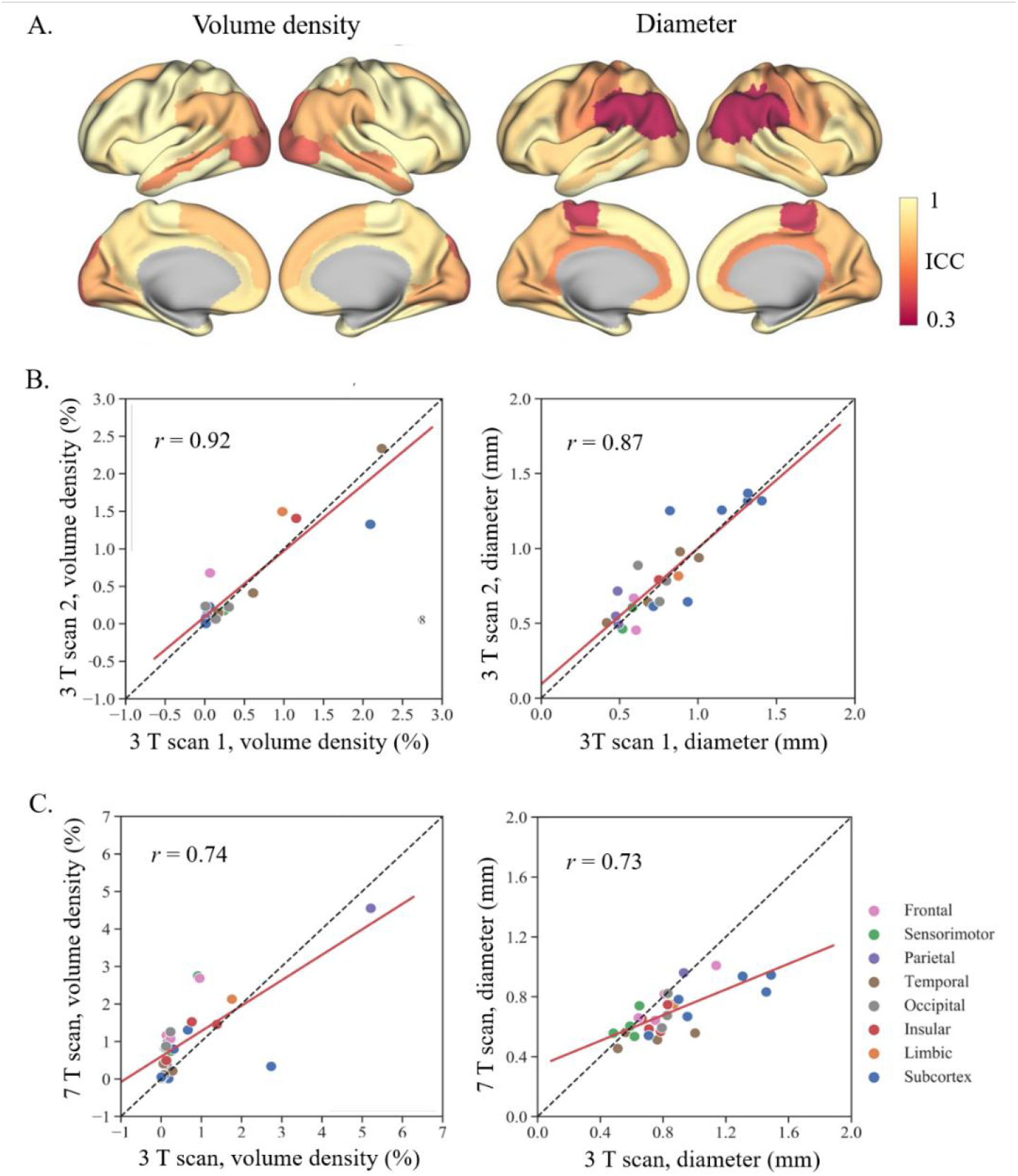
Test-retest results of region-specific volume density (left) and average diameter (right) for humans (A) and macaques (B, C). A, Intraclass correlation coefficient (ICC) overlaid on the human Brainnetome atlas for subjects in the MSC dataset. B and C, Interscan reliability based on the macaque Brainnetome atlas in two 3 T scans and in 3 T and 7 T scans in one macaque. The red line represents the best linear fit of the correlation between the measurements from two scans. The black dotted line represents a hypothetical complete agreement between the measurements from the two scans.

### 3.3 FFCM-MRF application

ROI-based and voxelwise group-averaged vascular volume density and diameter maps across the cortex for humans and macaques are shown in Figure 6 and Supplementary Figure 6. The patterns of vascular volume density and diameter were consistent across species, with dense and large arteries in the insular and anterior cingular regions and sparse and slender arteries in the lateral occipital region, indicating that the vascular pattern was highly conserved during evolution.

**Figure 6.**
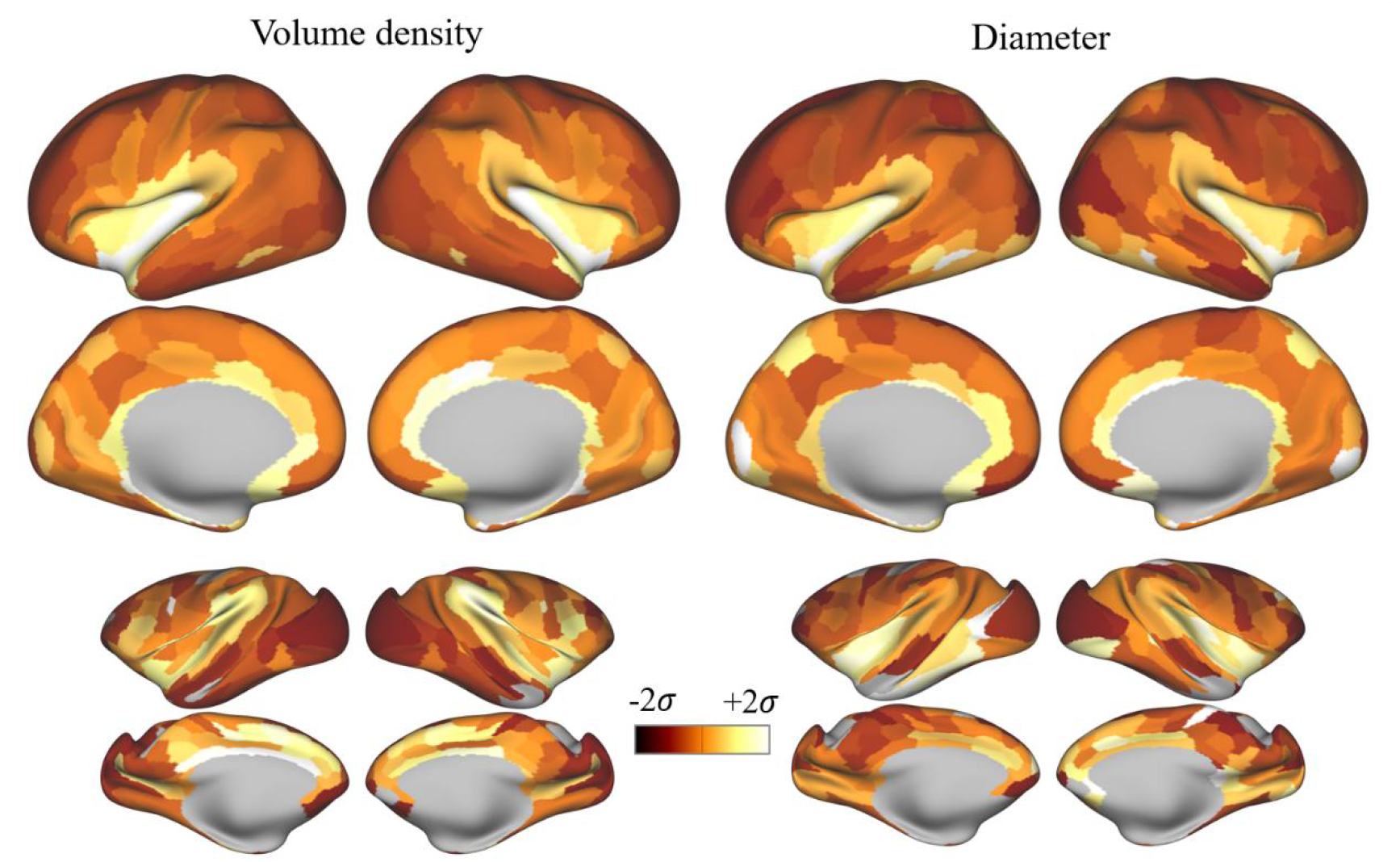
Group-averaged cerebrovascular volume density and diameter for humans and rhesus monkeys. Human and rhesus monkey Brainnetome atlases were used to extract the ROI-based vascular features.

## 4. Discussion

In the present study, we developed a fully automatic unsupervised cerebrovascular segmentation method, FFCM-MRF, which integrated fast FCM clustering with MRF while utilizing shape priors and spatial constraints. Compared with state-of-the-art deep learning and statistical model-based methods, this approach is highly accurate and reliable and largely independent of field strength, scanner type, acquisition parameters, and species.

Statistical cerebrovascular segmentation combined with MRF models has been reported previously (9–11). Most studies utilized finite mixture models based on the intensity distribution of TOF-MRA data, and an expectation maximization was performed to estimate the parameters. In contrast, this study used the FCM algorithm, which has shown superior performance for expectation maximization (12). In addition, we designed a novel neighborhood energy function with shape priors for MRF to reduce noise and improve segmentation performance. Compared with the GMM-MRF proposed by Zhang et al. (9), FFCM-MRF achieved an average increase of 3.21% and 4.91% in DSC across multiple human and macaque datasets, respectively, and was capable of effectively suppressing noise (Figure 2). Deep learning techniques have made rapid progress in cerebrovascular segmentation (3, 4, 20, 25). However, most cross-validation strategies were based on the same dataset (5, 6). A variety of scanners, acquisition protocols and species pose challenge for independent replications. We found that the proposed FFCM-MRF outperformed Transformer-based semi-supervised GCS models in all datasets with an average increase of 4.79% and 9.38% in DSC for humans and macaques, respectively. nnU-Net outperformed FFCM-MRF in the dataset for model training (i.e., MIDAS) with a higher DSC of 5.71%; however, FFCM-MRF improved the segmentation results in all the independent datasets by an increased DSC of 4.48% and 2.04% for humans and macaques, respectively (Table 3), suggesting that FFCM-MRF is more generalizable to new TOF-MRA images. The proposed method self-adaptively discriminates between vessel and non-vessel voxels based on intensity contrasts of individual images. As shown in Supplementary Figure 7, although the voxel intensity distributions were different between humans and monkeys, the vessel and non-vessel voxels exhibited well-isolated distributions in both species that could be identified and utilized for discrimination by FFCM. It is for this reason that vascular anatomical structural differences between humans and macaques (e.g., the lack of an anterior communicating artery in rhesus monkeys) do not affect segmentation results for rhesus monkeys. Rhesus monkeys are vital animal models in developing therapeutics for brain diseases. To the best of our knowledge, our work is first to generalize cerebrovascular segmentation algorithms to nonhuman primates. The effectiveness of FFCM-MRF for macaque cerebrovascular segmentation may promote progress in comparative neuroscience and translational medicine.

Because the vasculature annotations shown in Supplementary Figure 8 are voxel-based delineations on three views of 2D slices of TOF-MRA, the tubular and continuous shape information about the vessels was always cut into cross-sections. Without 3D intuitionistic vascular shape information, oversights always exist, especially for distal small arteries even if the annotator has experience and anatomical knowledge of the vascular system. Manual labeling of the whole brain’s sophisticated and complex vasculature network is therefore subjective, time consuming, and prone to inter-subject variability. In addition to generalizability, FFCM-MRF has unique superiority in detecting vessels that are ignored by annotators because the segmentation procedure is independent of ground truth labels. In contrast, deep learning models learn vascular representation by depending on large-scale high-quality ground truth labels, so the segmentation results may be compromised by inaccurate or insufficient annotations.

Qualitative comparisons between nnU-Net and FFCM-MRF showed that vessels near the edge of the brain/image, including major large vessels in the CW at the base of the brain and smaller distal arteries that cover the cerebral cortical surface, were often poorly detected by nnU-Net. Quantitative analyses of the CW arteries demonstrated that FFCM-MRF was more effective in segmenting the CW across all datasets, indicating that the proposed method is robust and superior for CW segmentation. Stenosis of the CW arteries restricts the passage of blood and significantly increases the risk of stroke (26), cognitive impairment (27, 28), and Alzheimer’s disease (29) and increases the risk of progression from mild cognitive impairment to dementia (30). For these reasons, accurate segmentation of the morphology of the CW is essential for early identification of cerebrovascular and neurodegenerative diseases.

Compared with 3 T MRA, high field strength 7 T MRA performed better in terms of signal-to-noise and contrast-to-noise ratio (31). Our model showed the best Dice scores for the 7 T human and macaque datasets compared with state-of-the-art methods and detected more slender vessels than for 3 T TOF-MRA. We observed that the segmentation accuracy for the 7 T Forrest dataset was poorer than for the other human 3 T datasets. This is possibly because the ground truth labels contained noise and compromised the Dice scores for this dataset (Figure 3A). Test-retest results showed that FFCM-MRF is robust and replicable with good intra-subject results in terms of arterial localization and structural morphology. Reliable and generalizable results suggest that the FFCM-MRF model has the potential to facilitate the use of an automated vascular identification system in real-world research and clinical environments with their heterogeneity in scanners and subjects.

In the process of brain development, vessels act as a powerful signaling system that mediates and guides cell migration, differentiation, and structural connectivity during the development of neurons and glial cells (32). This neurovascular interaction established during development continues to play a fundamental role in communication between the vascular and nervous systems throughout adulthood (33). Evidence has shown that morphological vascular features, such as volume and diameter, are interregionally heterogeneous across the brain (17, 18, 34) and are correlated with distinct neurons (34). Consistent with previous reports (17), the present study found various distributions of the volume density and lumen diameter of cortical arteries in humans, with dense and large cerebral arteries in the insular and anterior cingular regions and sparse and slender arteries in the lateral occipital area. Similar vascular patterns were found in rhesus monkeys in the present study. Previous findings have reported dense cerebrovascular networks in the primary sensory cortices in mice (34). However, we found that the primary sensory cortices had a medium vascular density in humans and macaques. This is possibly because the mice studies included the whole vasculature network including arteries, veins and capillaries using immunofluorescence staining or tissue transparency techniques. In contrast, in vivo TOF-MRA can image arteries and part of the veins at hundreds of micrometer resolution. Another explanation may be that there are dramatic differences in brain organization between rodents and primates. To our knowledge, this work is the first in which human and macaque vascular structures were extracted in vivo and showed that brain-specific vascularization distributions are highly conserved across primates. Accurate cerebrovascular segmentation for humans and macaques will provide benefits in understanding the evolution and pathophysiological underpinnings of diseases.

TOF-MRA allows for three-dimensional visualization and morphology quantification of the intracranial vessels in vivo and has become a powerful tool available in research and clinical settings. Robust, generalized and open-access tools, such as FFCM-MRF, have the potential to harmonize and process data from different sites, which represent real-world environments with their heterogeneity in participants and scanners. Although some semi-automated GUI tools have been developed, they are not publicly available (35), or they utilized traditional region-based/level set methods requiring heavy manual interventions (36). As of yet, there is no user-friendly open-source toolbox for automated cerebrovascular segmentation and analysis. The present study integrated FFCM-MRF segmentation with morphometric feature extraction into a user-friendly GUI toolbox (Supplementary files and a manual in the appendix) and is freely available at GitHub repository https://github.com/YueCui-Labs/FFCM-MRF. It was developed in MATLAB (MathWorks, Inc.) and includes TOF-MRA preprocessing, vascular segmentation, and feature extraction. Currently, there is also a lack of publicly available rhesus monkey angiography datasets. Macaque data from the Maca-7T and Maca-BJ datasets along with ground truth labels will be released on the OpenNeuro platform (https://openneuro.org) to promote the development of cutting-edge algorithms that can enable cerebrovascular segmentation and analysis across species.

## 5. Conclusions

This study proposed FFCM-MRF, an automated and open-source method for accurate intracranial cerebrovascular segmentation. The results demonstrated the superiority of our developed framework for brain vessel segmentation and its ability to generalize well to multiple human and macaque datasets. Because it can supply accurate regional features from segmentation, FFCM-MRF can facilitate the study of novel imaging biomarkers for cerebrovascular and neurodegenerative diseases, and thereby enable diagnoses and preventative treatments as early as possible.

## Data/Code availability

All TOF-MRA data used to develop and validate FFCM-MRF were acquired from different public human (MIDAS, BraVa, CHUV, ADAM, Forrest and MSC) and private macaque datasets (Maca-WH, Maca-BJ and Maca-7T). Macaque TOF-MRA data and ground truth labels from the Maca-BJ and Maca-7T have been released via OpenNeuro: https://openneuro.org/datasets/ds004620/versions/1.0.0. Data from the Maca-WH can be obtained upon reasonable request. URL links for public datasets are:

– MIDAS: https://public.kitware.com/Wiki/TubeTK/Data
– BraVa: http://cng.gmu.edu/brava/home.php
– CHUV: https://openneuro.org/datasets/ds003949/versions/1.0.1
– ADAM: https://adam.isi.uu.nl/
– Forrest: https://openneuro.org/datasets/ds000113/versions/1.3.0
– MSC: https://legacy.openfmri.org/dataset/ds000224/

FFCM-MRF pipeline GitLab repository: https://github.com/YueCui-Labs/FFCM-MRF.

## Author contributions

Guarantors of integrity of entire study, **Y.C., H.H., S.Y.**; study concepts/study design or data acquisition or data analysis/interpretation, all authors; manuscript drafting or manuscript revision for important intellectual content, all authors; approval of final version of submitted manuscript, all authors; agrees to ensure any questions related to the work are appropriately resolved, all authors; literature research, **Y.C., H.H., S.Y., M.Z., D.Y.**; experimental studies, **Y.C., H.H., J.L., C.L., M.Z.**; statistical analysis, **Y.C., H.H., J.L., C.L.**; and manuscript editing, all authors.

## Supplementary File

### Supplementary Figures

**Supplementary Figure 1.**
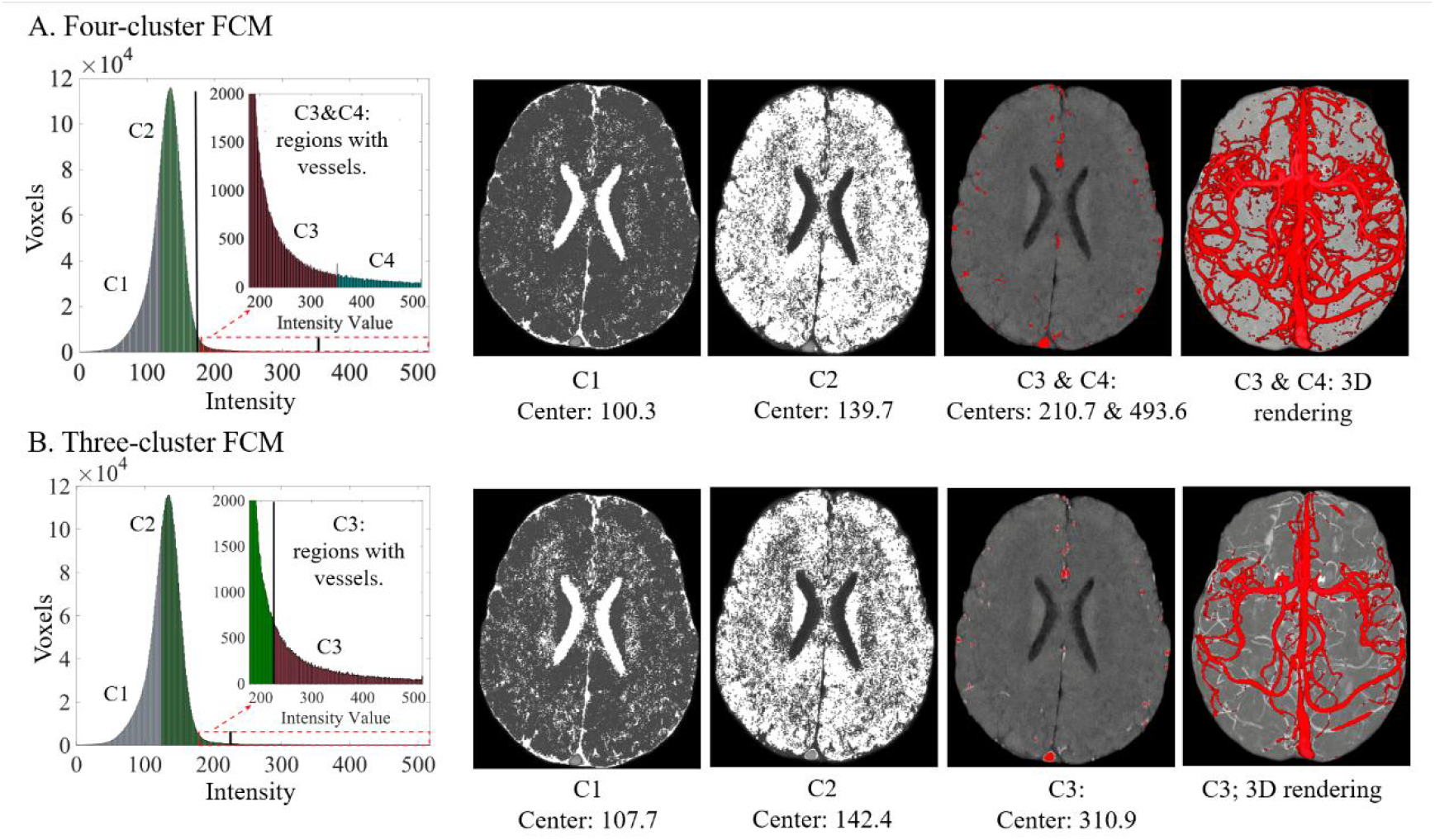
Results of three- (A) and four-cluster (B) fuzzy c-means (FCM) clustering on pre-processed time-of-flight magnetic resonance angiography (TOF-MRA). The first column shows the histograms colored by clustering results, and the second and third columns correspond to cerebrospinal fluid and brain tissues, respectively. The fourth column corresponds to cerebral vessels, and the fifth column represents the rendering of vessel segmentation overlaid on a maximum intensity projections view for the TOF-MRA image. The results demonstrate that cluster 3 (C3) and cluster 4 (C4) of the four-cluster FCM contained more cerebral vessels than cluster 3 (C3) of the three-cluster FCM.

**Supplementary Figure 2.**
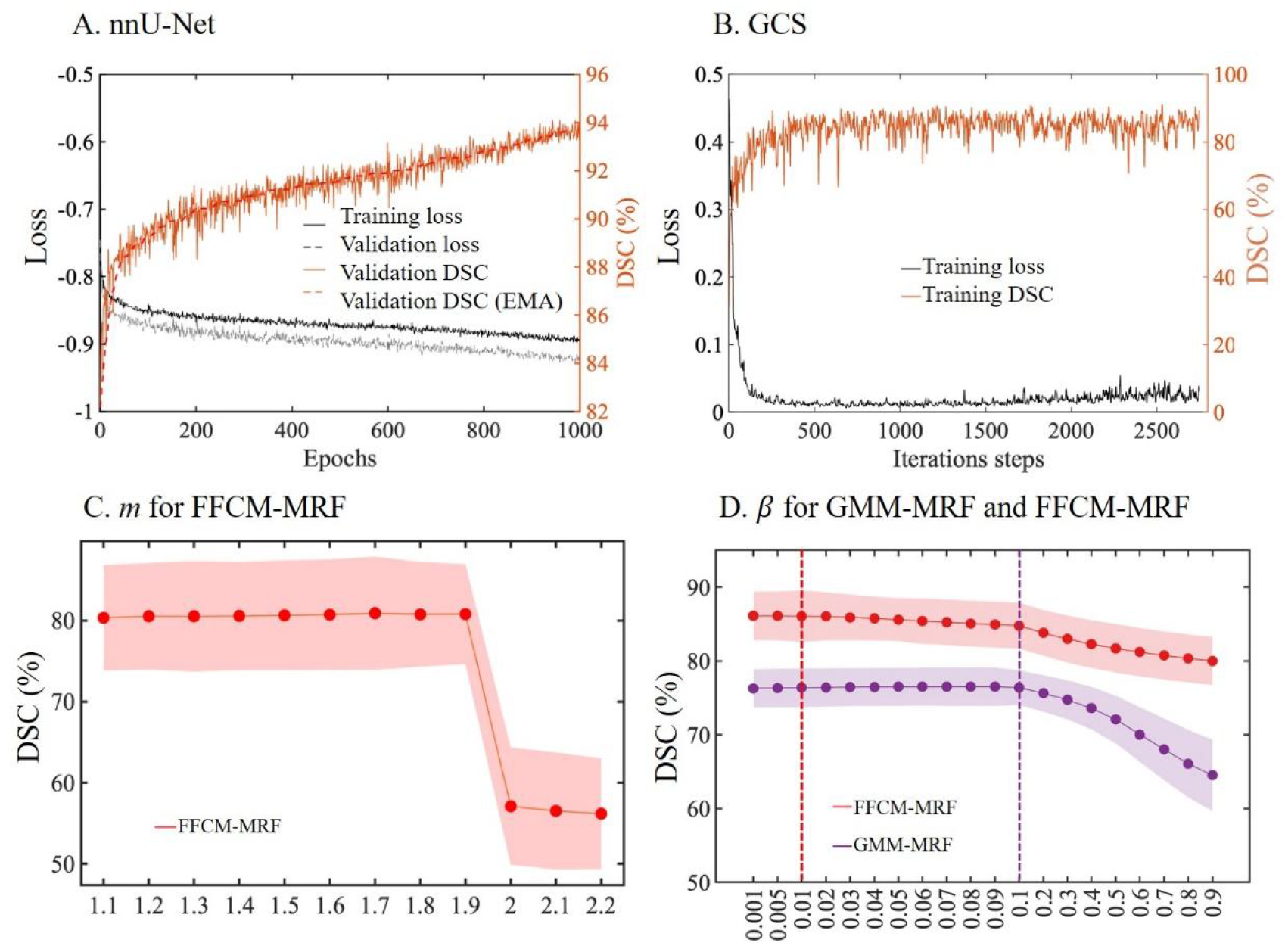
Model training procedures for nnU-Net and GCS, and hyperparameter tuning for GMM-MRF and FFCM-MRF methods in the MIDAS dataset. A, Model training procedure of nnU-Net. The model was trained for a total of 1000 epochs with 250 iterations per epoch. The graph shows the loss on the training and validation sets, the Dice score coefficient (DSC), and the exponential moving average (EMA) of the DSC score for the validation set. B, The loss and DSC score of each iteration of GCS on the training set were monitored. The GCS model was trained for 20 epochs with 135 iterations per epoch, resulting in a total of 2700 iterations as per (1). The best model from the training stage was used for the test and independent datasets. C, For FFCM-MRF, the optimal range of weighting exponent *m* for fuzzy c-means clustering was found to be between 1.1 and 1.9. D, The optimal β was set to 0.01 and 0.1 for FFCM-MRF and GMM-MRF, respectively.

**Supplementary Figure 3.**
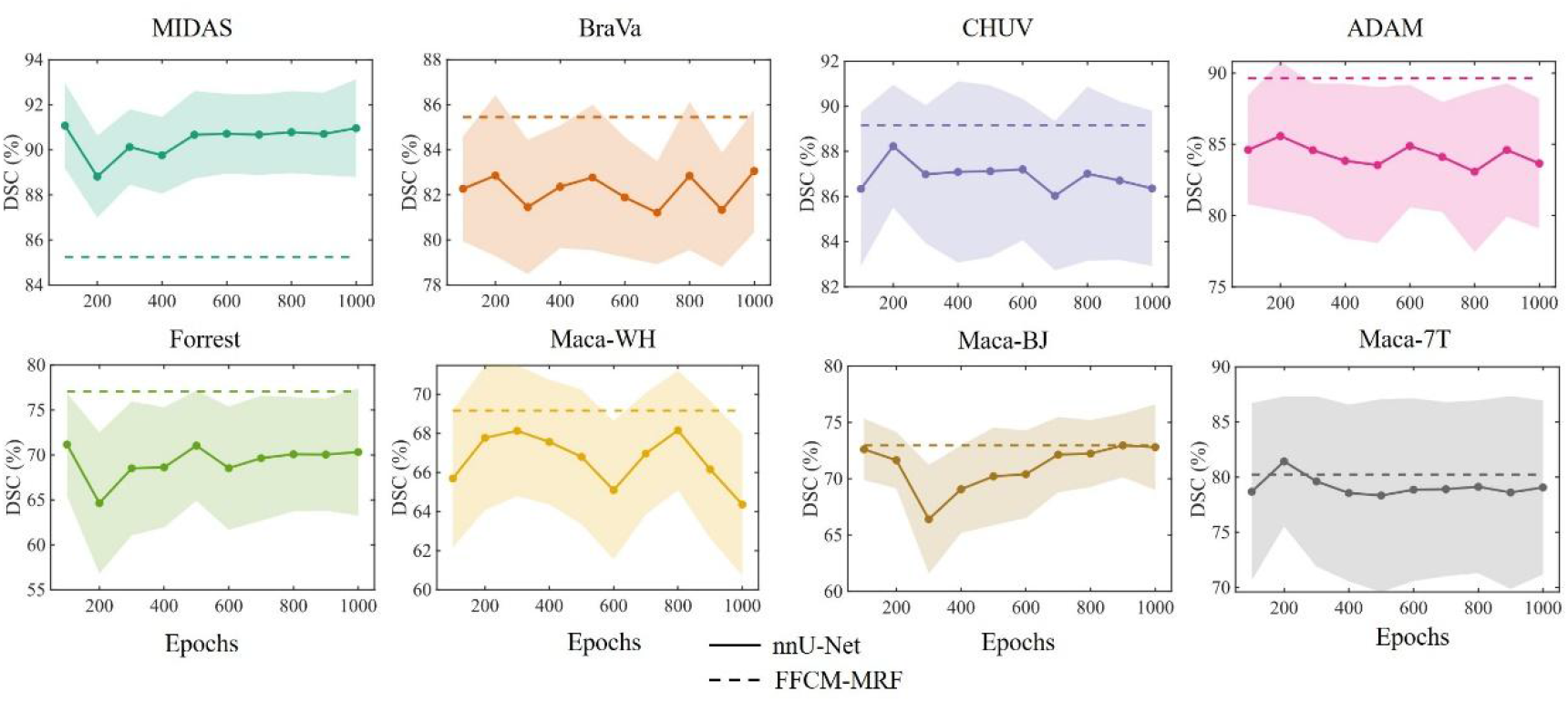
Experiments were conducted to evaluate the cerebrovascular segmentation performance using nnU-Net with 100 to 1000 epochs for different datasets. Our results showed that, except for MIDAS, FFCM-MRF outperformed nnU-Net within 1000 epochs in independent human and macaque datasets.

**Supplementary Figure 4.**
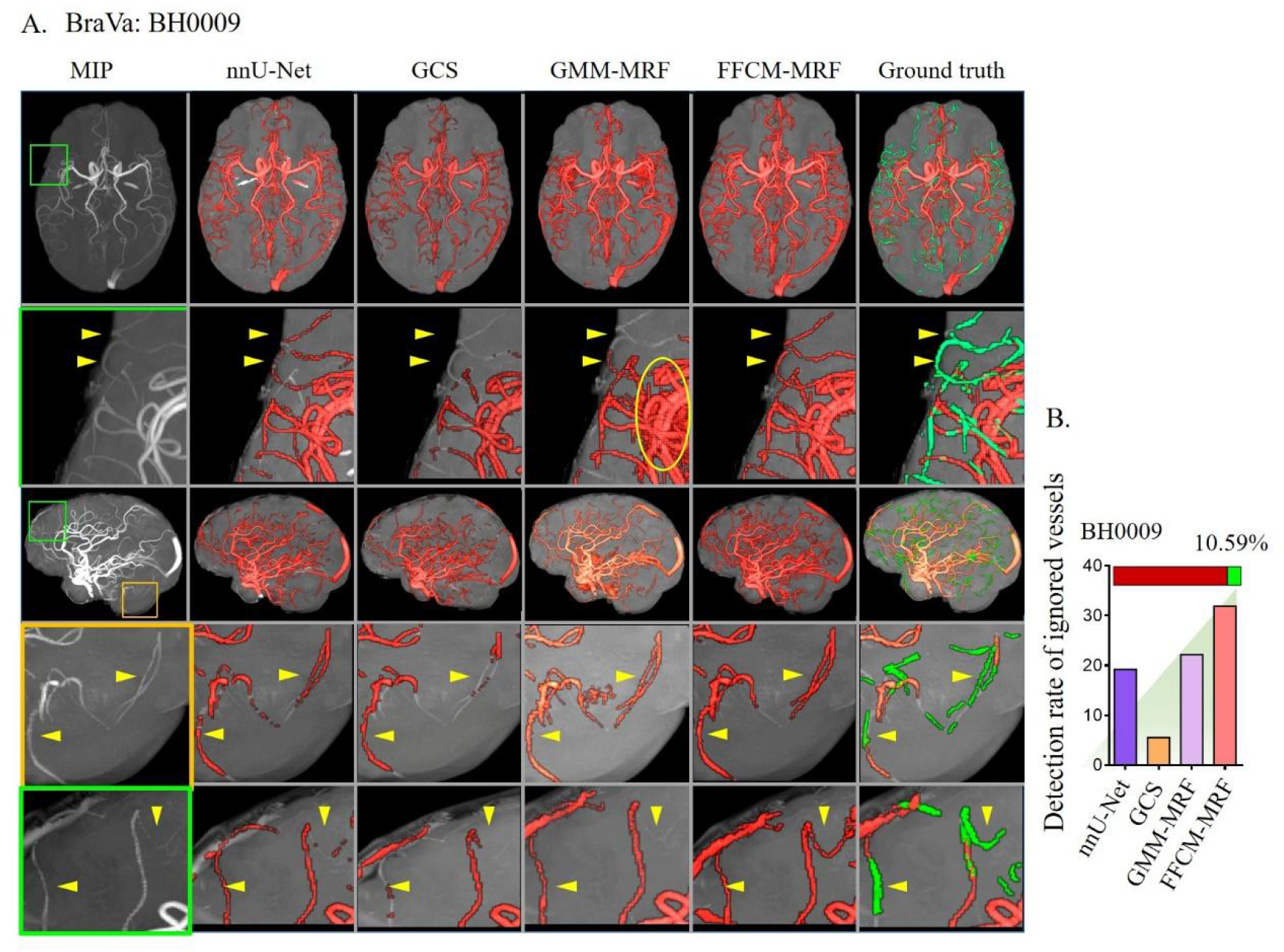
Comparisons of cerebrovascular segmentation results on BraVa dataset with state-of-the-art methods. A, Left to right columns correspond to maximum intensity projection (MIP) view for the TOF-MRA image, results by nnU-Net, GCS, GMM-MRF, and FFCM-MRF, the ground truth, and the ground truth with ignored vessels shown in green. The yellow arrows indicate that FFCM-MRF can detect vessels that were not annotated in the ground truth. The subject BH0009 was meticulously delineated by a trained annotator with the ignored vessels in green, as shown in the rightmost column in A. B, Detection rate of ignored vessels with state-of-the-art methods. Ignored vessels accounted for 10.59% of the whole brain vessels on TOF-MRA, indicated by the green (ignored) and red (ground truth) colors of horizontal bars in B.

**Supplementary Figure 5.**
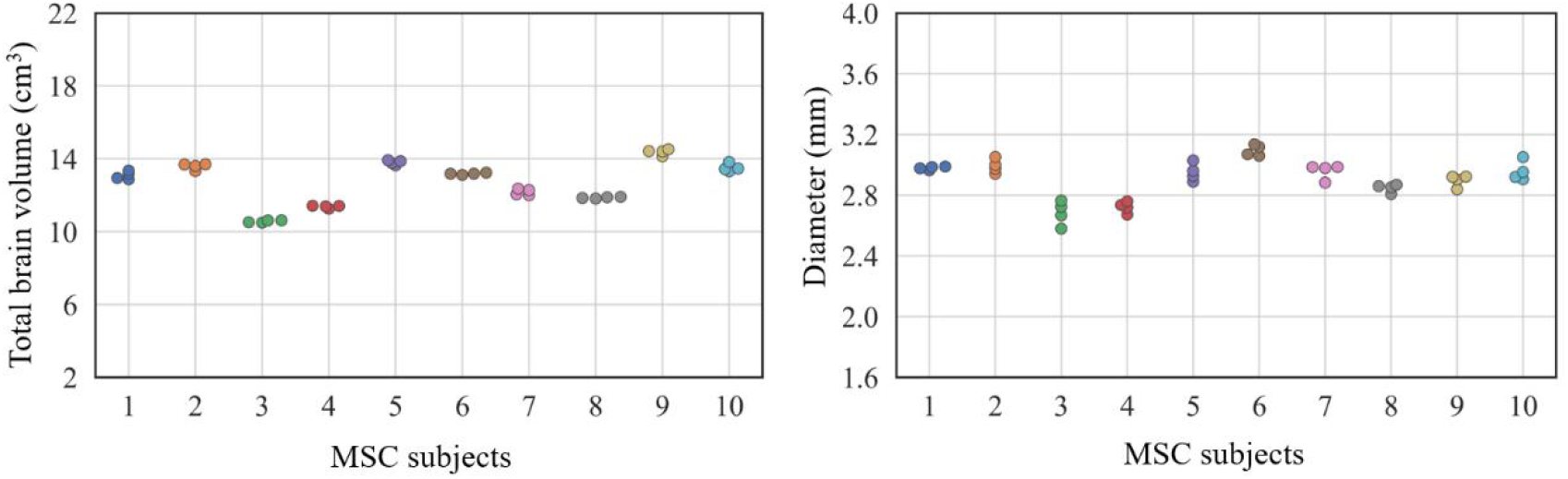
Test-retest results of total brain volume (left) and average diameter (right) from four scans in the MSC dataset.

**Supplementary Figure 6.**
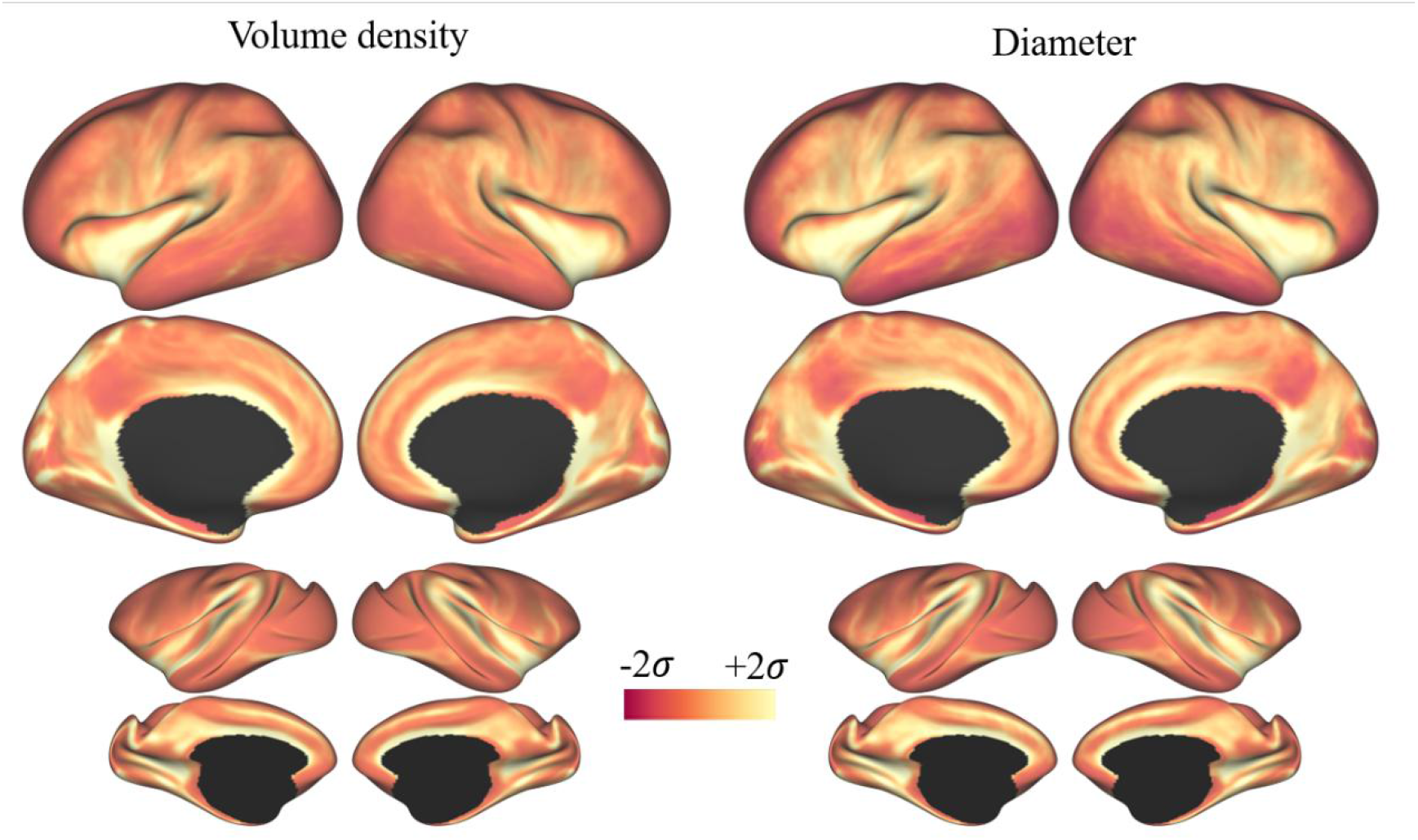
Group-averaged voxelwise cerebrovascular volume density and diameter for humans and macaque monkeys. These features were computed inside the bounding box of the 15-cube and 8-cube for humans and macaques, respectively. Each brain voxel served as the center of the bounding box, and the features within the bounding box were allocated to the voxel.

**Supplementary Figure 7.**
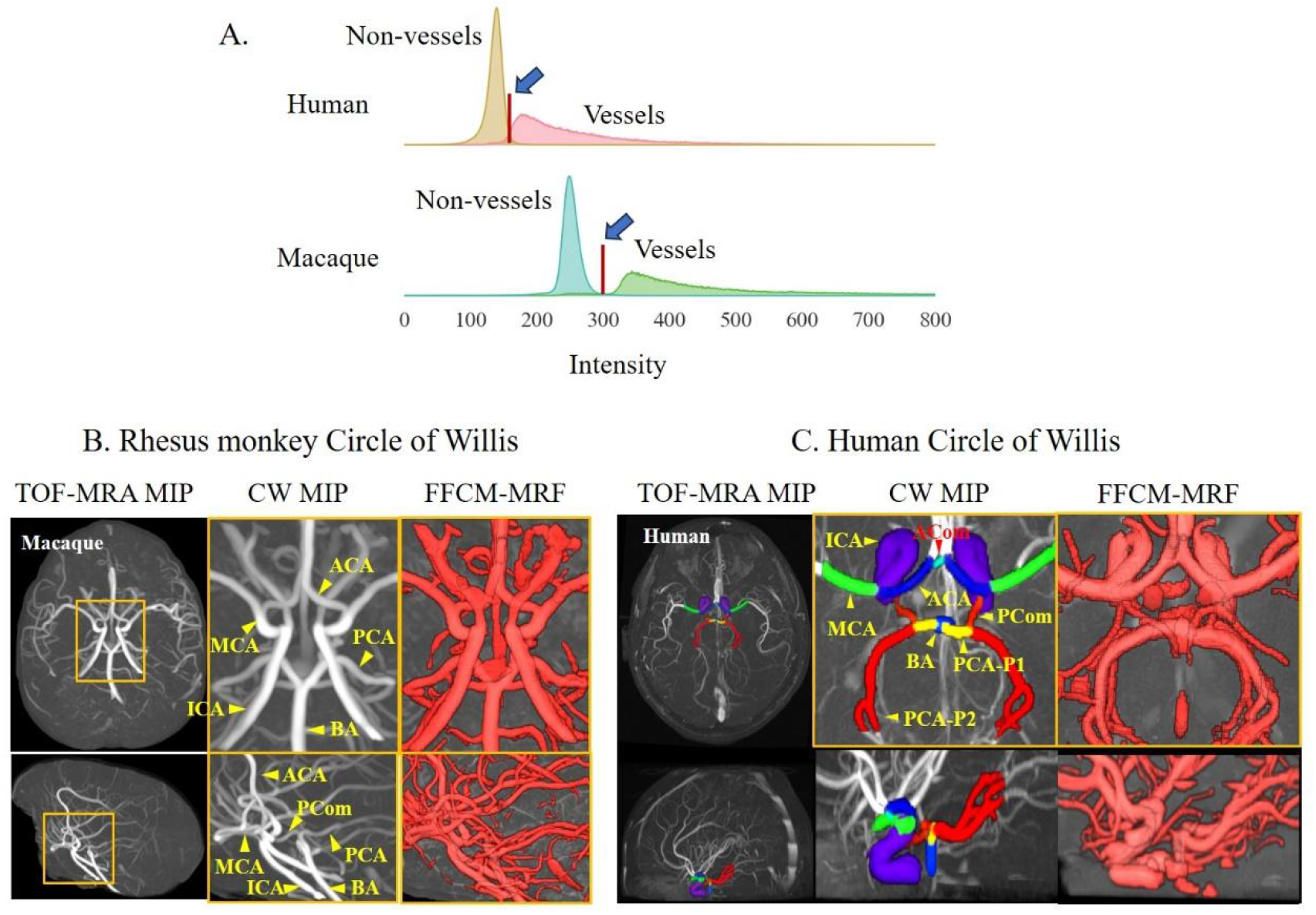
The histograms of vessel and non-vessel voxels for human and macaque subjects, and the Circle of Willis for rhesus monkeys and humans. A, Vessel and non-vessel voxels were plotted based on ground truth labels. Rudimentary segmentations were designed to adaptively identify the discrimination of intensity contrasts between vessel and non-vessel voxels on each individual image. Note that for visualization purposes, the vertical axes for vessel voxels have been expanded by approximately 200 times. B and C, The Circle of Willis for rhesus monkeys and humans. ACA, anterior cerebral artery; ACom, anterior communicating artery; BA, basilar artery; CW, Circle of Willis; ICA, internal carotid artery; MCA, middle cerebral artery; MIP, maximum intensity projection; PCA, posterior cerebral artery; PCom, posterior communicating artery.

**Supplementary Figure 8.**
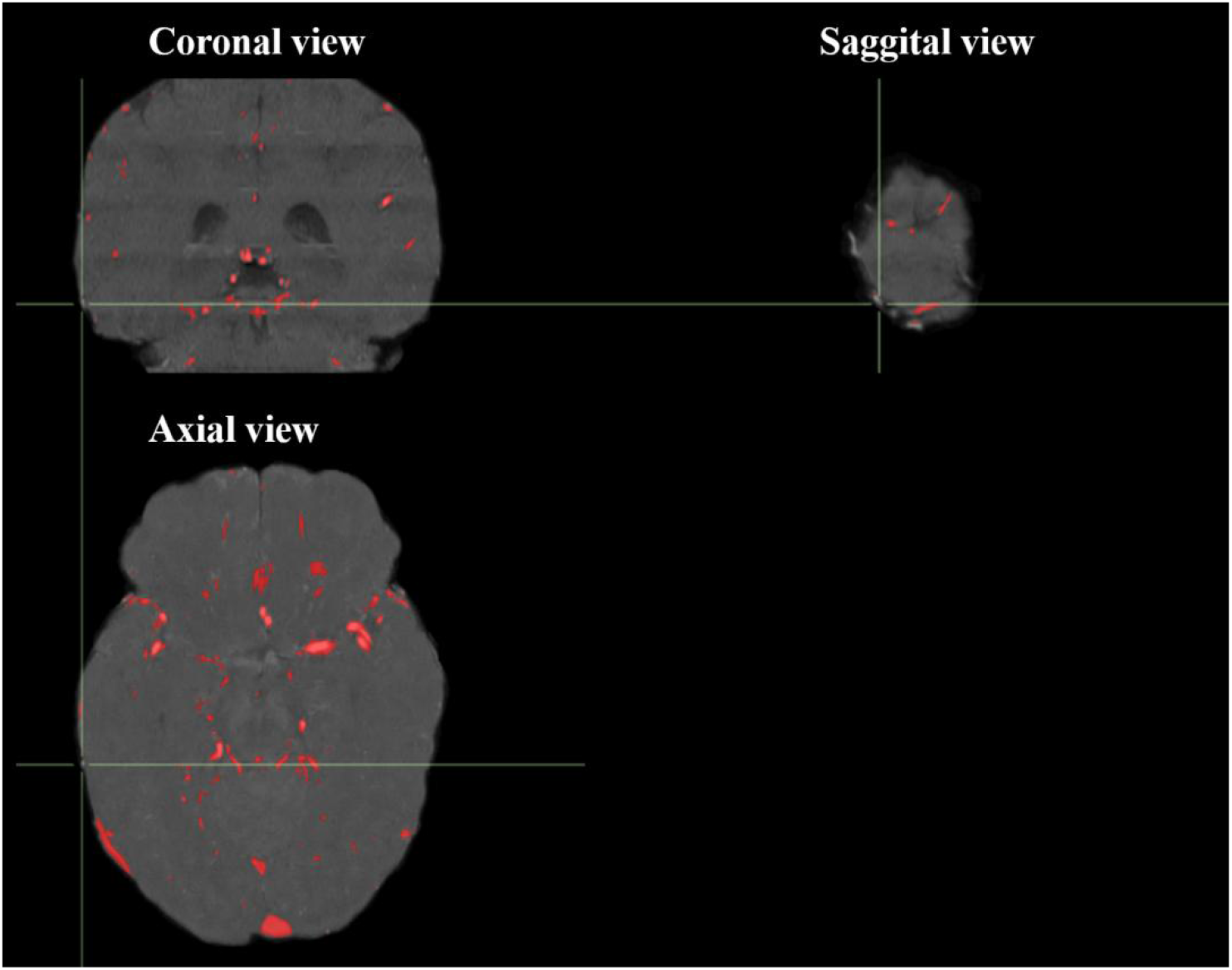
A TOF-MRA image was overlaid with ground truth labels. The location of the crossbar represents a segment of the pial artery that was not annotated by the ground truth labels.

**Supplementary Figure 9.**
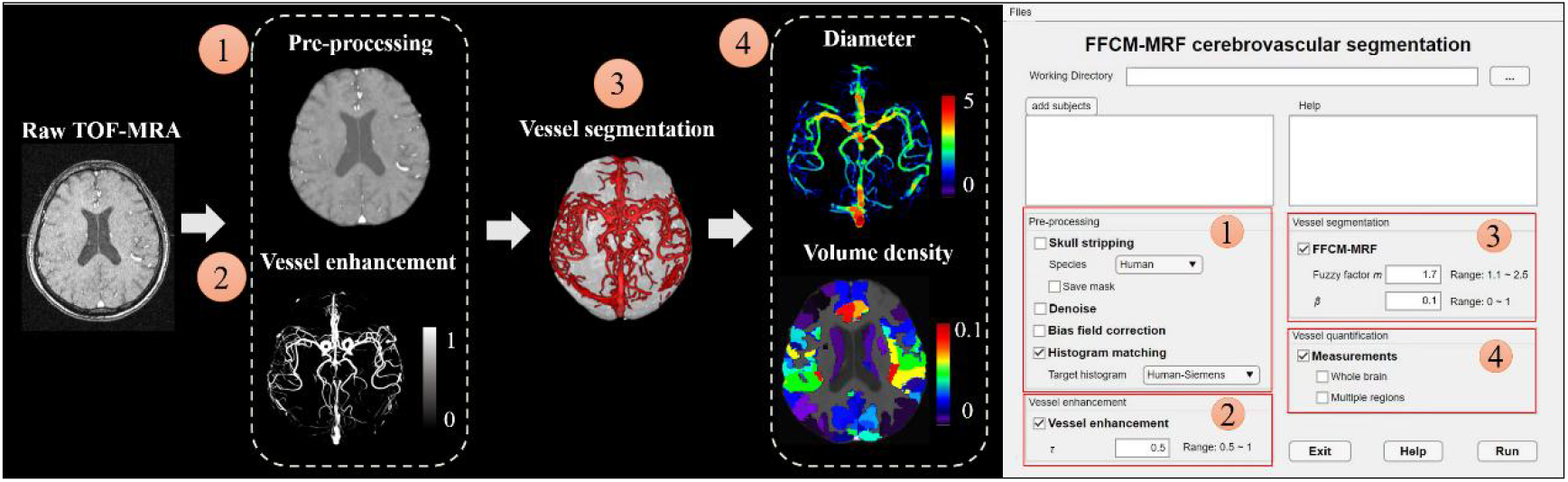
Graphical user interface and pipeline of FFCM-MRF.

**Supplementary Figure 10.**
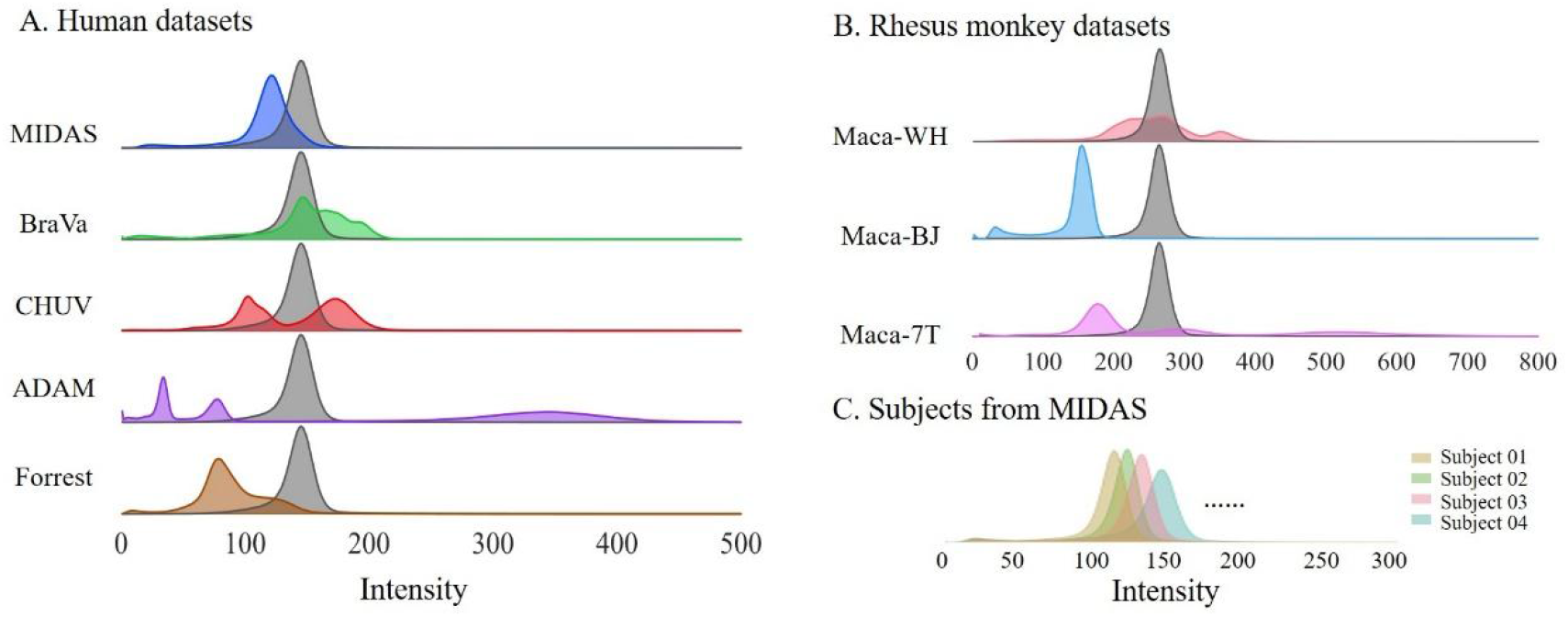
The histograms of skull-stripped TOF-MRA in multiple human and rhesus monkey datasets. A and B, The colored distribution represents the original histogram of each dataset. The gray shading behind each colored distribution represents the histogram after histogram specification. C, The histograms of different subjects from MIDAS dataset.

### Supplementary Tables

**Supplementary Table 1.** T1-weighted scans acquisition parameters. ^a^.

|  | Dataset | Scanner | Magnetic field strength (T) | Voxel resolution (mm<sup>3</sup>) | TR/TE (ms) | Flip angle (degree s) | Field of view (mm<sup>3</sup>) |
| --- | --- | --- | --- | --- | --- | --- | --- |
| Humans | MIDAS | Siemens | 3 | 1.00 × 1.00 × 1.00 | 1700/4.38 | 25 | 170 × 256 × 256 |
|  | BraVa | Siemens | 3 | 0.66 × 1.00 × 1.00 | 2100/3.04 | 11 | 256 × 256 × 256 |
|  | CHUV | Siemens Verio/<br>Philips Intera | 3 | 3.90 × 0.56 × 0.56 | 400/2.72 | 90 | 37 × 420 × 448 |
|  | Forrest | Philips Achieva | 3 | 0.7 × 0.67 × 0.67 | 2500/5.7 | 8 | 192 × 257 × 257 |
|  | MSC | Siemens Tim Trio | 3 | 0.80 × 0.80 × 0.80 | 2400/3.74 | 8 | 224 × 256 × 256 |
| Macaques | Maca-WH | Siemens Prisma | 3 | 0.50 × 0.50 × 0.50 | 2200/3.68 | 8 | 224 × 256 × 256 |
|  | Maca-BJ | Siemens Prisma | 3 | 0.50 × 0.50 × 0.50 | 2200/2.84 | 8 | 240 × 300 × 320 |
|  | Maca-7T | Siemens MAGNETOM | 7 | 0.50 × 0.50 × 0.50 | 2200/2.29 | 7 | 256 × 256 × 176 |

<sup>a</sup>T1-weighted scans were not accessible for ADAM dataset.

**Supplementary Table 2.**
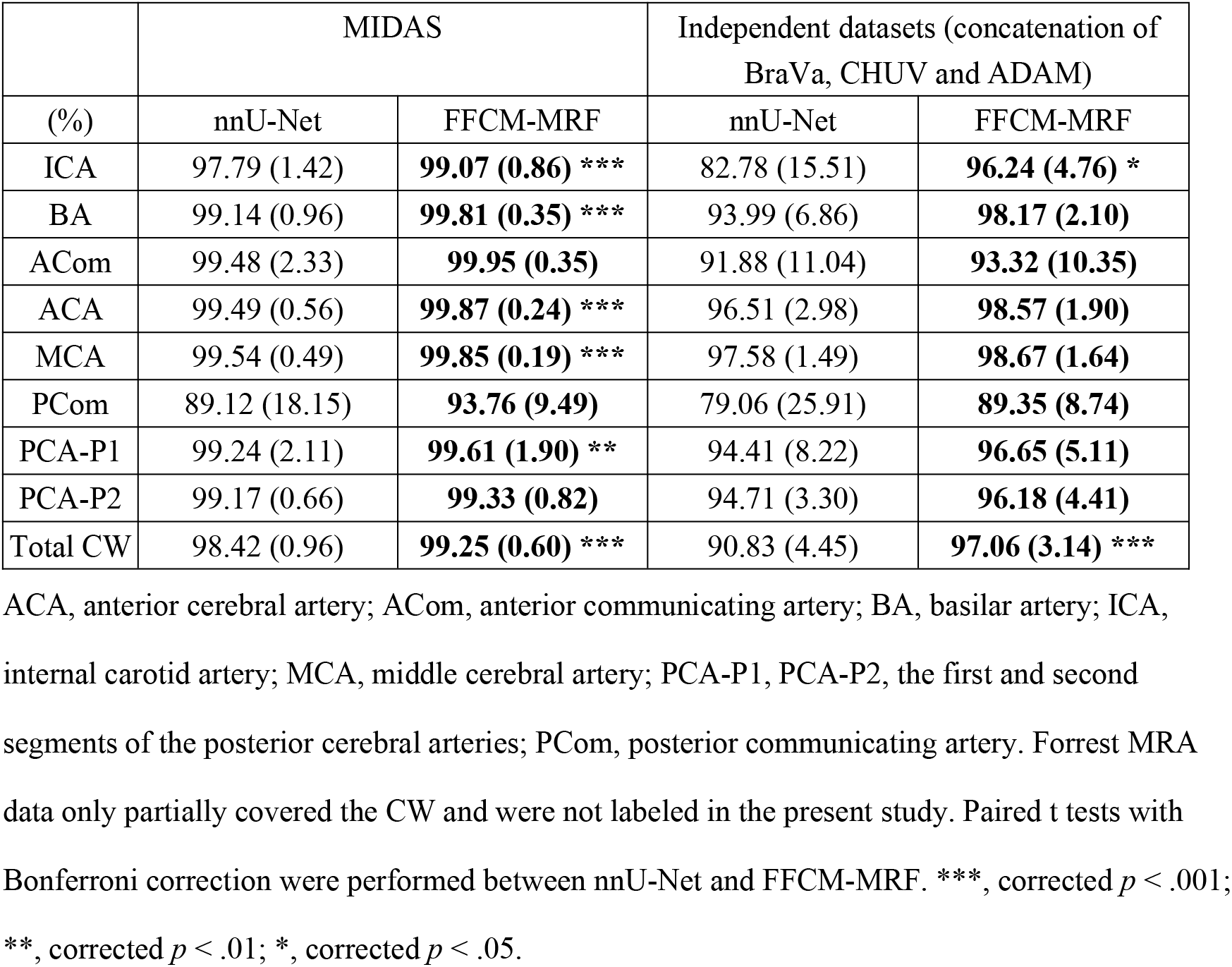
Detection rate for eight Circle of Willis arteries using nnU-Net and proposed FFCM-MRF methods.

**Supplementary Table 3.** Computation time of nnU-Net and FFCM-MRF per subject for different datasets.^a^.

|  |  |  | MIDAS | BraVa | CHUV | ADAM | Forrest |
| --- | --- | --- | --- | --- | --- | --- | --- |
| Intracranial voxels (k) |  |  | 6,813<br>(540) | 6,659<br>(539) | 8,221<br>(1,388) | 12,749<br>(753) | 30,501<br>(1,673) |
| Training/<br>parameter<br>tuning | nnU-Net (GPU) |  | 1083<br>(7.07) | — | — | — | — |
|  | FFCM-MRF (CPU) |  | 47.45<br>(7.41) | — | — | — | — |
| Inference/<br>segmentation | nnU-Net | CPU | 9.38<br>(0.02) | 11.03<br>(1.69) | 7.34<br>(1.35) | 6.84<br>(0.5) | 14.26<br>(0.17) |
|  |  | GPU<br>(seconds) | 8.35<br>(0.08) | 9.57<br>(1.54) | 6.22<br>(1.26) | 4.79<br>(1.48) | 2.93<br>(0.09) |
|  | FFCM-MRF<br>(CPU) | Histogram<br>specification | 1.15<br>(0.16) | 1.11<br>(0.15) | 2.00<br>(0.66) | 2.18<br>(0.54) | 2.52<br>(0.25) |
|  |  | Vessel<br>enhancement | 0.95<br>(0.04) | 0.79<br>(0.07) | 0.98<br>(0.19) | 1.31<br>(0.1) | 5.67<br>(0.24) |
|  |  | FFCM-MRF | 1.39 (0.1) | 1.22<br>(0.1) | 1.47<br>(0.22) | 2.23<br>(0.20) | 1.66<br>(0.10) |
|  |  | Total | 3.5 (0.1) | 3.11<br>(0.1) | 4.46<br>(0.35) | 5.73<br>(0.28) | 7.81<br>(0.40) |

<sup>a</sup> The unit of computation time is minutes unless otherwise indicated. Values are mean (SD).

**Supplementary Table 4.** Intraclass correlation coefficients of volume density and diameter based on the human Brainnetome atlas for the MSC dataset.

|  |  | Regions of interest | Volume density<br>ICC | Diameter ICC |
| --- | --- | --- | --- | --- |
| 1 | Frontal Lobe | SFG, Superior Frontal Gyrus | 0.75 | 0.91 |
| 2 |  | MFG, Middle Frontal Gyrus | 0.98 | 0.83 |
| 3 |  | IFG, Inferior Frontal Gyrus | 0.90 | 0.93 |
| 4 |  | OrG, Orbital Gyrus | 0.86 | 0.85 |
| 5 |  | PrG, Precentral Gyrus | 0.97 | 0.70 |
| 6 |  | PCL, Paracentral Lobule | 0.96 | 0.39 |
| 7 | Temporal Lobe | STG, Superior Temporal Gyrus | 0.96 | 0.94 |
| 8 |  | MTG, Middle Temporal Gyrus | 0.63 | 0.80 |
| 9 |  | ITG, Inferior Temporal Gyrus | 1.0 | 1.0 |
| 10 |  | FuG, Fusiform Gyrus | 0.93 | 0.82 |
| 11 |  | PhG, Parahippocampal Gyrus | 0.95 | 0.96 |
| 12 |  | pSTS, Posterior Superior<br>Temporal Sulcus | 0.88 | 0.90 |
| 13 | Parietal Lobe | SPL, Superior Parietal Lobule | 0.97 | 0.78 |
| 14 |  | IPL, Inferior Parietal Lobule | 0.74 | 0.33 |
| 15 |  | Pcun, Precuneus | 0.85 | 0.76 |
| 16 |  | PoG, Postcentral Gyrus | 0.94 | 0.61 |
| 17 | Insular Lobe | INS, Insular Gyrus | 0.97 | 0.82 |
| 18 | Limbic Lobe | CG, Cingulate Gyrus | 0.90 | 0.63 |
| 19 | Occipital Lobe | MVOcC, MedioVentral Occipital<br>Cortex | 0.71 | 0.73 |
| 20 |  | LOcC, Lateral Occipital Cortex | 0.51 | 0.81 |
| 21 | Subcortical Nuclei | Amyg, Amygdala | 0.96 | 0.79 |
| 22 |  | Hipp, Hippocampus | 0.99 | 0.89 |
| 23 |  | BG, Basal Ganglia | 0.87 | 0.70 |
| 24 |  | Tha, Thalamus | 0.98 | 0.97 |

### Supplementary materials and methods

#### 1. MRA datasets

A total of 123 human and 44 macaque MRA images scanned at 3 T and 7 T MRI from 9 datasets was used to develop and validate FFCM-MRF. Human MRA data were taken from the publicly available MIDAS (2), BraVa (http://cng.gmu.edu/brava/home.php), CHUV (3) (https://openneuro.org/datasets/ds003949/versions/1.0.1), ADAM (https://adam.isi.uu.nl), Forrest (4) (https://www.studyforrest.org), and Midnight Scan Club (MSC, https://legacy.openfmri.org/dataset/ds000224/) datasets. The MIDAS database was released by the CASILab at the University of North Carolina at Chapel Hill. The present study included 54 MRA images with voxel-wise segmentation ground truth labels, of which 20 were manually labeled by Hilbert et al. (5), and 34 were released by the CASILab (https://public.kitware.com/Wiki/TubeTK/Data). Note that the segmentation ground truth labels only focused on arteries, but we also included veins, such as the venous sinus, because they are prominent vessels in MRA images. Subsets of ADAM, BraVa, and CHUV were used as independent datasets to validate the FFCM-MRF. The Forrest dataset comprises 7 T high field strength high-resolution MRA images from 20 participants. Of these images, 14 were labeled by SMILE-UHURA Challenge 2023 (https://www.soumick.com/en/uhura/). Forrest MRA acquisition was performed with partial brain coverage and high resolution. Test-retest validation was performed based on the MSC dataset, which included four repeated TOF images for each of 10 participants.

36 Macaque MRA images were acquired from a 3 T Siemens Prisma scanner in Wuhan (Maca-WH), 4 were acquired from a 3 T Siemens Prisma scanner from the Institute of Biophysics, Chinese Academy of Sciences in Beijing (Maca-BJ), and 4 were obtained from a 7 T MAGNETOM Terra scanner (Maca-7T) at the same institute. The monkeys were anesthetized with an intramuscular injection of Zoletil 50 (10 mg/kg) before scanning and were maintained under anesthesia using 0.5% isoflurane in oxygen during image acquisition (6). The animals were placed in a sphinx position during the scan, with an MRI-compatible stereotaxic frame optimized for the NHP head. Ground truth labels for ADAM, BraVa, CHUV, and macaques were labeled by a trained annotator and two students (authors H.H. and J.L.). See Tables 1 and 2 for subject details and references for acquisition parameters. This study was approved by the Institutional Review Board/Ethics Committee of Chinese Academy of Sciences Institute of Automation.

#### 2. MRA pre-processing

The MRA images were first skull stripped using the *bet* function in FSL 6.0 for the humans, and DeepBet (7) for the macaques to remove non-brain tissues, followed by *DenoiseImage* and *N4BiasFieldCorrection* functions in ANTS for denoising and bias field correction. The extracted brains were carefully reviewed in case manual edits were required using ITK-SNAP (http://www.itksnap.org/pmwiki/pmwiki.php). To mitigate the intensity variability across scanners and subjects, we used histogram specification to normalize these images in order that data from different sites are uniformly distributed (Supplementary Figure 10). Specifically, we first calculated the peak value of the histogram for each subject in the MIDAS dataset, and then identified the peak value that was most common among the subjects, and finally averaged the histograms with that peak value to obtain the target histogram. The histograms of all the subjects in MIDAS, BraVa, CHUV, ADAM, and Forrest were then matched to the target histogram established using MIDAS. For macaques, the target histogram was generated for the Maca-WH dataset, and then all the macaque MRA images were aligned to it.

#### 3. The FFCM-MRF segmentation procedure is illustrated in Algorithm 1

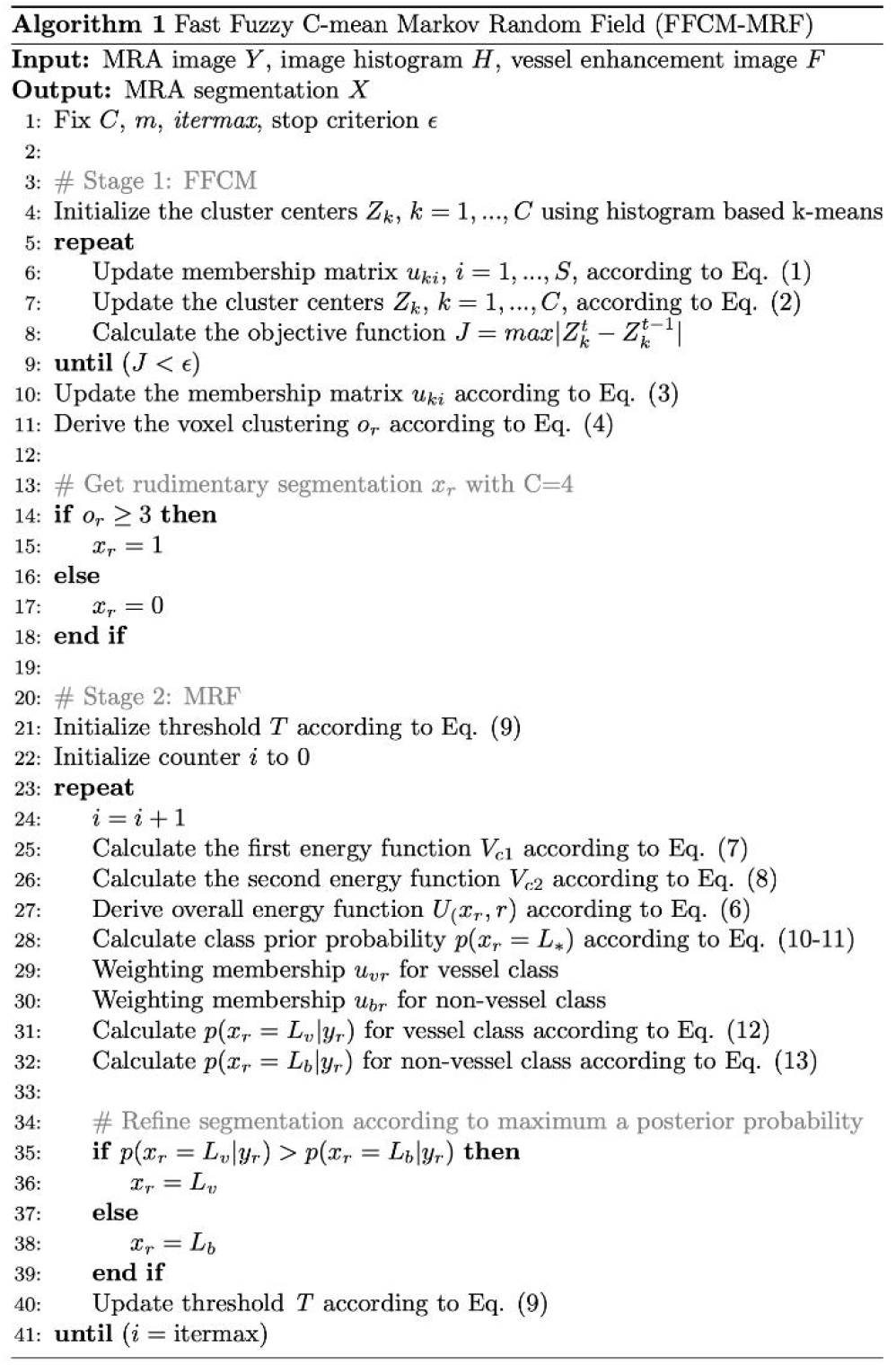

#### 4. FFCM-MRF quantitative evaluation

95HD (95% Hausdorff distance): The Hausdorff distance (HD) measures the distance between two binary volumes, i.e., prediction and ground truth. HD is defined as

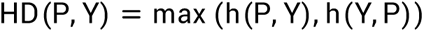

where h(P, Y) is called the directed Hausdorff distance and is given by

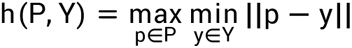

where ||p-y|| is the Euclidean distance between points p and y. 95 percentile Hausdorff distance (95HD) is commonly used to avoid outliers.

AHD (Average Hausdorff distance): The Average Hausdorff distance (AHD) is the HD averaged over all voxels. The AHD is known to be stable and less sensitive to outliers than the HD or 95HD.

The AHD is defined as

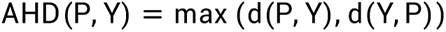

where d(P, Y) is the directed average Hausdorff distance that is given by

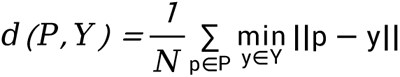

### 5. Comparisons with other state-of-the-art models

FFCM-MRF was compared with two state-of-the-art deep neural network models, nnU-Net (8) and GCS (1), as well as with a statistical model, GMM-MRF (9). The input images for these three models were pre-processed TOF-MRA images. nnU-Net is an ensemble of U-Net architecture with an automated pipeline and is currently state-of-the-art for training biomedical segmentation tasks. We optimized nnU-Net for 100 epochs with a batch size of 2 and 250 batches per epoch. Ablation experiments were performed with epochs from 100 to 1000. The batch size was optimized as 160 × 192 × 80, with at least one-third of the patches guaranteed to contain vessels to ensure robust handling of class imbalances. nnU-Net resamples all images at the same target spacing by default. Since the MRA images for the macaques had a relatively high resolution compared with those for humans, resampling was not performed for the macaques to avoid down-sampling, which may potentially result in a poor segmentation performance.

GCS is a Transformer-based semi-supervised cerebrovascular segmentation method, which utilizes reconstruction consistency to constrain segmentation models and improve the texture representation of the model. Because GCS is a semi-supervised method, an additional 182 unlabeled images from multiple datasets (MIDAS n = 55, BraVa n = 51, CHUV n = 55, ADAM n = 15 and Forrest n = 6) were used for the GCS model training. All hyper-parameters remain consistent with those in Chen et al. (1). These included the random cropping strategy, which was applied to crop the TOF-MRA to a size of 128 × 128 × 128, and the maximum epoch was set as 20. For each epoch, every image was cropped into 10 patches, which formed 135 batches and each batch contained 2 patches from a labeled image and 2 patches from an unlabeled image.

To ensure generalization and reduce variability, a 2-fold cross-validation was employed using the MIDAS dataset for training nnU-Net and GCS. This involved randomly partitioning the MIDAS dataset into two subsets, with one of these used as test data and the remaining one as training data. This procedure was repeated twice, with each subset used as test data once. The final segmentation results for MIDAS represented the average of these two independent experiments, and the model that performed better on the test set was considered to be the optimal model and was used for independent validation of all the other datasets. For nnU-Net and GCS, the models were all implemented using the Pytorch framework (v2.0.0) and trained on a high-performance server using a single NVIDIA GeForce RTX 3090 GPU. For GMM-MRF, we selected the same participants from MIDAS with FFCM-MRF to tune parameter *β* by maximizing the DSC, and the optimal *β* was then used for all MIDAS and other datasets. Models for GMM-MRF and FFCM-MRF were implemented using MATLAB2017b on a PC Intel(R) Xeon(R) Gold 5318Y CPU @2.10 GHz.

